# “Recurrent Extinction of Resistance Mutations Leads to Convergent Multidrug Resistance in Sequential Antibiotic Treatment”

**DOI:** 10.1101/2025.04.08.647880

**Authors:** Adam Lyon, Muhammed Sadik Yildiz, Samuel Cooke, Scott H. Saunders, Adam Rosenthal, Erdal Toprak

**Affiliations:** Department of Pharmacology, The University of Texas Southwestern Medical Center, Dallas, TX 75390, USA; Department of Microbiology and Immunology, University of North Carolina, Chapel Hill, NC, USA; Lyda Hill Department of Bioinformatics, The University of Texas Southwestern Medical Center, Dallas, TX 75390, USA

## Abstract

To overcome the antibiotic resistance problem, exploiting reproducible evolutionary tradeoffs is considered key for designing evolution-proof antibiotic therapies. Yet the predictability of resistance evolution has been viewed as limited, given the often idiosyncratic genetic trajectories observed in laboratory evolution experiments. To address this, we partially mimicked clinical antibiotic pharmacodynamics by imposing strong selection and evolved *Escherichia coli* under single or sequential antibiotic treatment. Under single-antibiotic selection, endpoint resistance, persistence, and tolerance phenotypes were reproducible, but populations evolving in parallel frequently followed divergent genetic trajectories. Remarkably, sequential antibiotic use redirected these divergent paths toward genotypic and phenotypic convergence, driven by extinction of resistance-conferring mutations when switching to the next effective antibiotic. Single-cell RNA sequencing demonstrated that evolved cultures contain cells occupying distinct metabolic niches and include more cells in states with lower translational activity and higher expression of toxin-antitoxin genes. These findings provide evolutionary insights to inform clinically effective antibiotic treatment strategies that employ sequential treatment.

## INTRODUCTION

Antibiotic resistance is a global health crisis causing over one million deaths annually^1,2^. With limited new antibiotics in development, evidence-based antibiotic choices are essential to prolong the efficacy of current and future antibiotics. Patients requiring prophylaxis or those with serious infections undergo multiple rounds of antibiotic treatment. Unfortunately, these complex treatment regimens are often established haphazardly with little guidance for clinicians. Repeated treatment failures can lead to multi-drug resistant (MDR) infections, which are associated with higher morbidity and mortality^3^. Therefore, antibiotic choices must consider both clinical efficacy and the evolutionary consequences of treatment failures to mitigate these risks.

Laboratory evolution experiments have revealed many insights about canonical resistance mechanisms such as reduced drug uptake, increased efflux, and drug inactivation. However, we now know there are many other non-canonical antibiotic survival strategies, and these conventional experimental evolution protocols may not capture these non-canonical mechanisms because they do not account for the pharmacological complexity of drug treatment in patients^4–6^. In clinical settings, antibiotic concentrations routinely peak and trough, consistently remaining above the minimum inhibitory concentration (MIC), a clinical metric used to determine susceptibility of a pathogen to a given antibiotic (Figure 1a)^7^.

**Figure 1.**
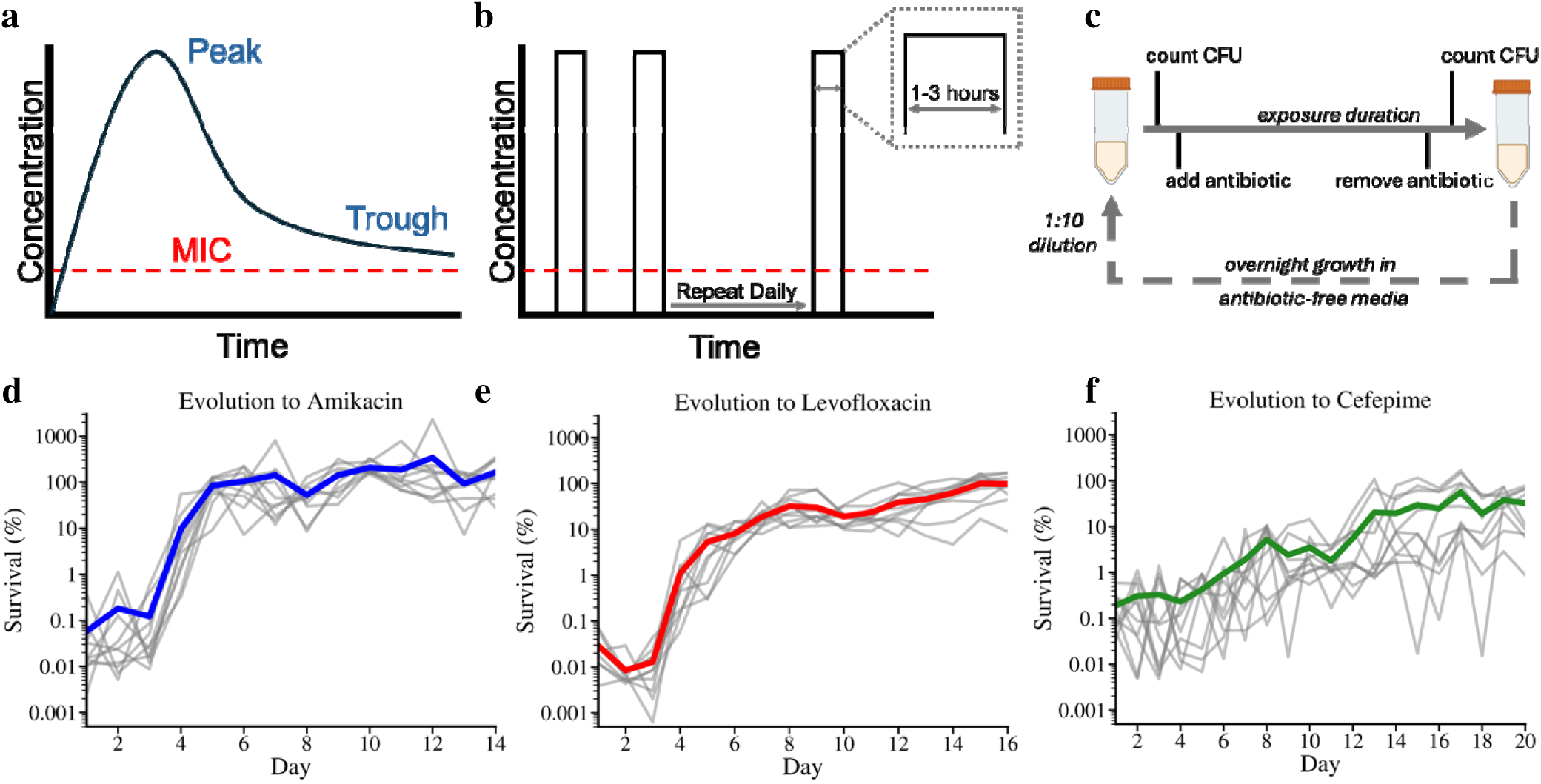
Evolution protocol rationale, scheme, and survival outcomes of single antibiotic evolutions. **a,** Illustration of the pharmacokinetics in a clinical antibiotic regimen with peak and trough antibiotic concentrations remaining above the minimum inhibitory concentration (MIC). **b,** Laboratory adaptation of clinical parameters for evolution experiments featuring daily antibiotic exposures at peak concentrations with adjustable exposure durations. c, Complete evolution protocol workflow. CFU counts are measured before and after antibiotic addition and removal, followed by overnight growth in antibiotic-free media and a 1:10 dilution after overnight growth. d-f, Survival trajectories of bacterial cultures evolved in the presence of amikacin, levofloxacin, and cefepime, respectively. Gray lines represent individual replicate cultures, while colored lines represent the mean survival over time for amikacin (blue), levofloxacin (red), cefepime (green).

Antibiotic persistence and tolerance are two non-canonical bacterial survival mechanisms about which we have limited knowledge^8–10^. In antibiotic tolerance (Supplementary Figure 1), the entire bacterial population survives longer under antibiotic exposure due to a slower killing rate, despite having the same MIC as antibiotic-sensitive bacteria^11^. Persistence, by contrast, arises from a subpopulation of bacteria that survives antibiotic exposure by dying more slowly, leading to a biphasic killing curve (Supplementary Figure 1). These slower killing rates pose significant challenges for treatments aiming to completely eradicate infections. However, our understanding of tolerance and persistence remains limited, as they are difficult to study^12^. Moreover, their true prevalence in clinical settings is unknown because they are often miscategorized as antibiotic susceptible due to indistinguishable MIC values.

To study the evolution of resistance, persistence, and tolerance under the selection of three clinically important antibiotic compounds, we utilized a laboratory evolution protocol that partially mimics antibiotic pharmacodynamics in clinical treatments by using strong selection while maintaining population diversity. Using this protocol, we evolved *Escherichia coli* populations with single or multiple, sequential antibiotics until they achieved sustained survival. Although phenotypic outcomes were similar, selection with single or sequential antibiotics drove distinct and reproducible differences in the associated mutational trajectories. We found that bacteria employ diverse resistance, persistence, and tolerance mechanisms, specific to each antibiotic, to evade strong killing effects induced by antibiotics. Most strikingly, sequential antibiotic use shifted evolutionary patterns from divergence to convergence, driven by recurrent extinction of resistance-conferring mutations. Despite convergent genetic trajectories in populations evolved under sequential antibiotic use, single-cell RNA sequencing of selected cultures revealed both unique and shared gene expression patterns across distinct single-cell clusters.

## RESULTS

We used three antibiotic compounds throughout this work: levofloxacin, amikacin, and cefepime. Each compound represents a separate antibiotic class, and they were chosen due to their frequent empirical and prophylactic use in clinics, particularly in pediatric stem cell transplant patients^13–15^. Levofloxacin is a fluoroquinolone that targets DNA synthesis through DNA gyrase inhibition^16^. Amikacin is an aminoglycoside that binds the 30s ribosomal subunit causing mistranslation which in turn kills bacteria through proteotoxic stress^17,18^. Cefepime is a cephalosporin, a beta lactam antibiotic, that disrupts a key step in cell wall synthesis^19^.

To quantitatively evaluate the evolutionary consequences of prolonged single antibiotic treatment, we evolved ten replicate isogenic cultures of a laboratory strain of *E. coli* (K-12, MG1655, abbreviated here as MG) to levofloxacin, amikacin, or cefepime (30 cultures in total). In addition, four untreated isogenic MG cultures were passaged daily without any antibiotic exposure as a control.

Our evolution protocol was designed to reveal clinically relevant bacterial survival strategies and associated genetic changes. Our protocol involves daily exposures of an antibiotic at a dose corresponding to its reported peak serum concentration in human patients, with durations optimized to achieve approximately 99.9% initial killing of the sensitive parent bacteria (Figure 1b). Here, the doses we used were 8.1 µg/ml of levofloxacin, 69 µg/ml of amikacin, and 129 µg/ml of cefepime^20–22^. These doses are higher than the MIC values of the parent strain (Levofloxacin: ∼0.11 µg/ml; Amikacin: ∼47.08 µg/ml; Cefepime: ∼0.18 µg/ml, Figure 2b-d, Supplemental Figure 2).

**Figure 2.**
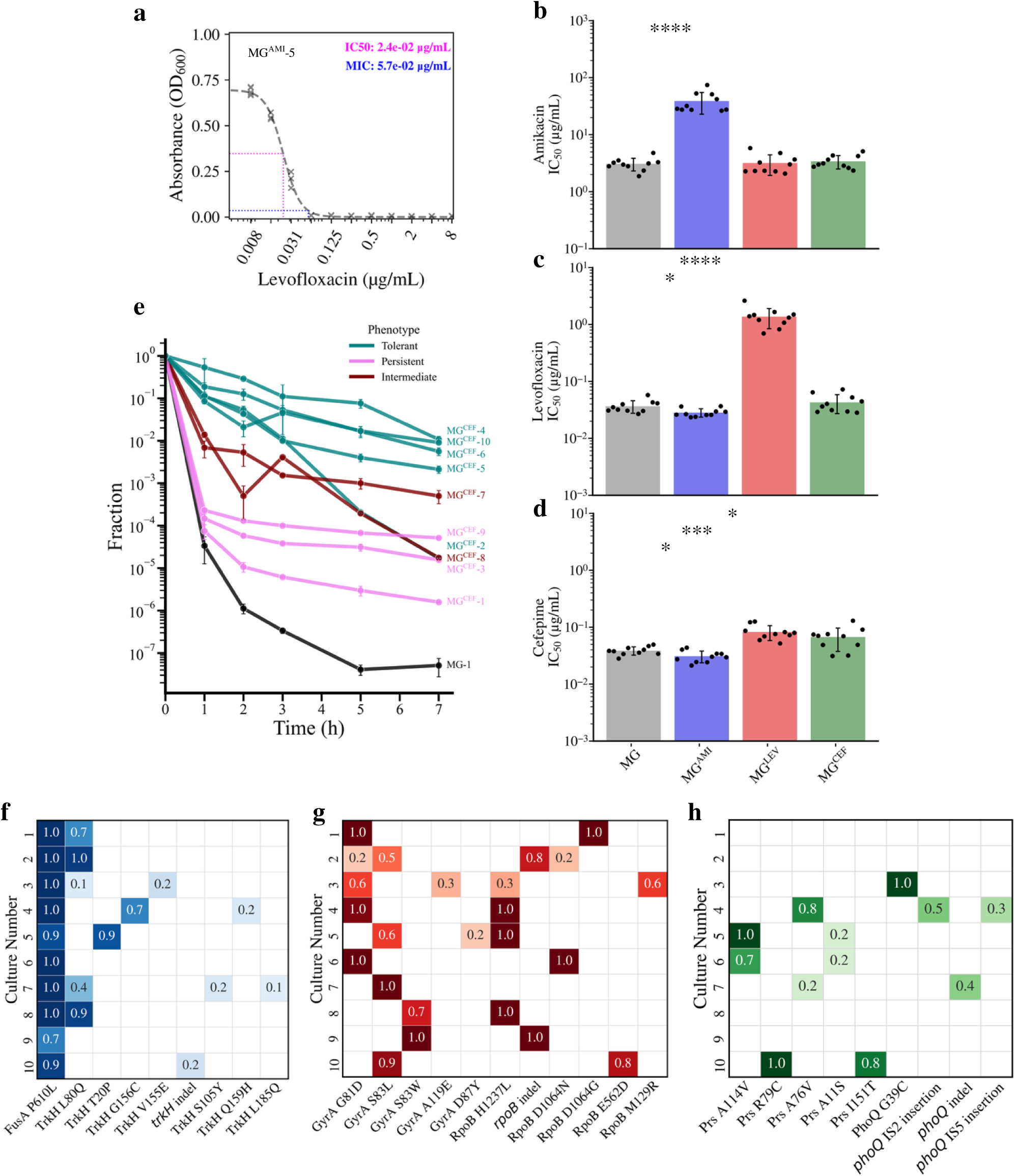
Phenotypic and genotypic outcomes of evolutions to single antibiotics. a, representative IC_50_ and MIC determination from a dose-response curve. b-d, bar plots of IC_50_ values in antibiotics amikacin, levofloxacin, and cefepime, respectively for parental and evolved cohorts (MG: gray; MG^AMI^: blue; MG^CEF^: green; MG^LEV^: red). Bar height corresponds to the mean IC_50_, and error bars reflect the standard deviation. Individual IC_50_ values for replicate cultures are shown as dots. Statistical significance (p-value, student’s t-test) is denoted by asterisk (* <0.05; **<0.01, *** < 0.001, **** < 0.0001), a convention used in subsequent figures. e, MDK assay for cefepime-evolved cultures (MG^CEF^) showing the median survivor fraction plotted over time, with error bars representing the median absolute deviation (MAD). Line colors differentiate the parental strain (MG: black) and evolved cultures that are characteristically tolerant (teal), persistent (pink), or of an intermediate phenotype (maroon). f-h, heatmaps of selected mutations and their frequencies among individual cultures: f, MG^AMI^ (blue); g, MG^LEV^ (red); g, MG^CEF^ (green).

Throughout the evolution experiments, we sampled the populations daily to quantify bacterial survival by counting colony forming units (CFUs) before and after antibiotic exposure. Antibiotics were removed by washing, and the surviving bacterial populations were transferred to antibiotic-free growth media (LB: Lysogeny Broth, all experiments were performed at 37 °C) for overnight growth. The next day, a modest 10-fold dilution and outgrowth primed the population for the next antibiotic exposure to avoid significant population bottlenecks and loss of diversity which are common in laboratory evolution protocols^23,24^ (Figure 1c). This protocol was repeated over several weeks until bacterial populations largely survived (>30 percent) daily antibiotic exposure as measured by percent survival calculated from CFU counts before and after treatment.

The initial killing was strong for each antibiotic. Average survival on Day 1 to levofloxacin, amikacin, and cefepime was 0.029 ± 0.02 %, 0.058 ± 0.11 %, and 0.527 ± 0.49%, respectively (mean ± standard deviation; n = 10; Figure 1d-f). As the daily selection cycles continued, levofloxacin and amikacin treatments led to survival trajectories that rapidly increased with a sigmoidal shape toward robust average survivals of 97.2 ± 51.8% and 163.7 ± 116.9% (mean ± standard deviation) after 16 and 14 treatment days, respectively (Figure 1d-e). Cefepime treatment also ultimately led to a treatment failure equivalent. However, unlike survival trajectories for populations evolved under the selection of levofloxacin or amikacin where the trajectories had sigmoidal shapes, those observed in cefepime treatment gradually increased toward a muted survival with a mean survival of 33.8 ± 25.1% despite 20 treatment days (Figure 1f). There was also more daily variation in survival among the individual trajectories relative to what was observed in experiments under levofloxacin or amikacin selection.

### Amikacin resistance and collateral sensitivity evolve through *fusA* and *trkH* mutations

Next, we assessed if increased bacterial survival was linked to known antibiotic resistance mechanisms. We generated dose response curves for each culture of evolved and parent cohorts in all three antibiotics to quantify cross resistance or collateral sensitivity (Figure 2a). Both MIC and inhibitory concentrations that reduced bacterial growth by half (IC_50_) were determined (Supplemental Figure 2, Methods). We then employed population Illumina whole genome sequencing (Methods) of bulk cultures, both evolved and parent, to identify mutations and their frequencies within and among the replicate evolved populations.

Cultures evolved under amikacin selection (MG^AMI^) gained highly significant resistance toward amikacin with the average IC_50_ increasing ∼12.7-fold (Figure 2b, p = 7.8×10^-5^, student’s t-test). Modest but statistically significant collateral sensitivities^5,25,26^ (reductions in IC_50_) were observed toward levofloxacin (Figure 2c, p = 0.012) and cefepime (Figure 2d, p = 0.037).

All ten MG^AMI^ cultures strikingly shared a single high frequency mutation of *fusA* (P610L), a gene which encodes ribosomal elongation factor G that processes the translocation of the ribosome during translation and has been associated with aminoglycoside resistance^5,25^. Additionally, *trkH*, a gene encoding a potassium ion channel^27,28^, was a common mutational target. Eight of ten cultures had at least one of eight unique TrkH mutations: L80Q, T20P, G156C, V155E, S105Y, Q159H, L185Q, or a 9-nucleotide deletion causing a frameshift after codon 385. TrKH L80Q was the most common mutation and present in five of ten cultures (Figure 2f). TrkH mutations in aminoglycoside resistant bacteria and their association with collateral sensitivity were previously reported by several groups including ours^25,29^. Additional disparate mutations were observed among the replicate cultures (Supplemental Figure 5a).

### Levofloxacin resistance is associated with *gyrA* and *rpoB* mutations

Cultures evolved under levofloxacin selection (MG^LEV^) acquired resistance toward levofloxacin with the average IC_50_ increasing ∼37.5-fold (Figure 2c, p = 2.4×10^-5^, student’s t-test). No significant change in amikacin susceptibility was observed in MG^LEV^ populations (Figure 2b). However, mild cross resistance to cefepime was observed (Figure 2d, p = 7.0×10^-4^).

All MG^LEV^ cultures had a mutation of *gyrA* which encodes DNA gyrase, a target of levofloxacin, consistent with previous reports^30^. Five unique GyrA mutations were observed: G81D, S83L, S83W, A119E, and D87Y, ordered by highest occurrence (Figure 2g). In addition to GyrA mutations, six MG^LEV^ cultures had a synonymous mutation of isocitrate dehydrogenase^31–33^ (*icd*: Icd H366H, cac➔ cat), and nine cultures had a mutation of RNA polymerase beta subunit^34^ (*rpoB*: RpoB H1237L, indel, D1064N, D1064G, E562D, and M129R). Culture 2 had both RpoB D1064N and the *rpoB* indel with different frequencies (Figure 2g). Although *rpoB* mutations have been reported following ciprofloxacin selection, the mechanism underlying this association remains unknown^34^. Interestingly, we have not observed any efflux related mutations in MG^LEV^ cultures even though such mutations are common in cultures evolved in the presence of gradually increased levofloxacin or ciprofloxacin concentrations^5^.

### Evolution with cefepime promotes tolerance and persistence, with divergent genetic changes

To our surprise, cultures evolved under cefepime selection (MG^CEF^) failed to produce robust resistance to cefepime, despite surviving treatment at cefepime doses almost a thousand-fold higher than the MIC. The IC_50_ of cefepime rose only modestly (<2-fold) (p= 0.014) (Figure 2d). Additionally, no significant cross resistance was observed toward amikacin (Figure 2b, p = 0.288) or levofloxacin (Figure 2c, p = 0.324).

We next asked whether bacterial survival in the presence of an extremely high cefepime concentration was due to a noncanonical survival mechanism such as antibiotic tolerance or persistence. Minimum duration for killing (MDK) measurements were previously proposed to identify and quantitatively differentiate these phenotypes (Supplementary Figure 1)^11^. To perform these measurements, cultures undergoing a continuous high-dose antibiotic treatment are periodically sampled for survival (measured by CFU counts) to determine the surviving fraction relative to the untreated cultures. Over 7 hours, we monitored the killing effects experienced by the parent strain (MG) and cefepime evolved (MG^CEF^) cultures (Figure 2e). Three MG^CEF^ cultures (numbers 1, 3, and 9) exhibited a biphasic shift in the killing effect at 1 hour in response to the treatment indicative of a persistence phenotype (Figure 2e, pink lines). A cohort of five MG^CEF^ cultures (numbers 2, 4-6, 10) exhibited a characteristically tolerant phenotype (Figure 2e, teal lines) with a steady and slower killing effect relative to the parent strain. Two MG^CEF^ cultures (numbers 7 and 8) have a biphasic switch at the 1-hour time point indicative of persistence, but the survivor fraction was much larger than the previously described persistent cohort. With one-hour survivor fractions of ∼10^-2^, these cultures appear to have a larger persistent subpopulation or an intermediate phenotype with characteristics of both persistence and tolerance (Figure 2e, maroon lines).

To compare our persistent and tolerant populations to bona fide resistance, we evolved cefepime resistant *E. coli* populations (MG^CEF-R^) using a conventional laboratory evolution protocol and evolved six replicate isogenic cultures of MG under cefepime selection. We grew the cultures in a dilution series of cefepime concentrations, and each day, for 15 days, we propagated the cultures surviving in the highest concentrations allowing growth into increasingly higher concentrations of antibiotic (Supplemental Figure 3a). This resulted in significant resistance to cefepime with the average IC_50_ rising ∼227-fold (p = 0.027) (Supplemental Figure 3d). MG^CEF-R^ cultures also evolved cross resistance to levofloxacin (Supplemental Figure 3c), but limited change in amikacin susceptibility (Supplemental Figure 3b).

We also tested whether this protocol-related effect could be generalized beyond our laboratory strain and if the outcomes could be considered clinically relevant. Previously in our lab, a clinical, non-pathogenic *E. coli* isolate (PEc) was genetically barcoded (PbEc, abbreviated here as Pb) and used to evaluate the evolution of resistance and persistence to cefepime in mice^5^. In that study, *in vitro* evolution assays using the conventional protocol were also conducted and produced bona fide cefepime resistance (Pb^CEF-R^). We evolved six cultures of Pb under the selection of cefepime using our present protocol for 15 days (Pb^CEF^). Like its MG^CEF^ counterpart, we found that the survival trajectory of Pb^CEF^ was gradual toward muted survival (Supplemental Fig 4a), and ultimately there was no increase in cefepime resistance (Supplemental Fig 4d). No changes in IC_50_ were observed for Pb^CEF^ populations against amikacin or levofloxacin (Supplemental Figure 4b-c). Instead, MDK assays revealed distinct tolerant and persistent cohorts among evolved populations (Pb^CEF^, Supplemental Fig 4e).

Genetic trajectories of cultures evolved under cefepime selection (MG^CEF^) were highly divergent, resulting in numerous unique mutations and idiosyncratic genetic changes (Figure 2h). Mutations in MG^CEF^ cultures were predominantly found at low frequencies (∼10-25%, Supplemental Figure 5c) and only a few mutations were present in more than one culture. Two genes, *prs* and *phoQ*, were common mutational targets among subsets of MG^CEF^ cultures. *prs*, encoding PRPP synthase (ribose-phosphate pyrophosphokinase), had five unique mutations (A114V, R79C, A76V, A11S, and I151T) present in five of the ten MG^CEF^ cultures, with no single mutation occurring in more than two cultures (Figure 2h). Likewise, *phoQ*, encoding a transmembrane histidine kinase, had four unique mutations (G39C, an indel, and two different mobile insertion elements) each present in only one MG^CEF^ culture (Figure 2h). Mutations of *prs* correspond to MG^CEF^ cultures observed as having a tolerant phenotype in the MDK assay (Figure 2e, teal), consistent with previous reports^35^. No pattern between mutations of *phoQ* and a survival phenotype (e.g. persistence) was apparent.

In contrast, cefepime resistant MG^CEF-R^ populations had convergent evolution toward mutations in genes known to be associated with beta lactam resistance including peptidoglycan transpeptidase^36^ (*ftsI*) and with multidrug resistance including outer membrane porin C^37^ (*ompC*), an *ompC* repressor, *envZ*^36,38^, efflux pump gene *acrB* of the *acrAB* operon, its repressor *acrR*^36,39^, and the *marR* repressor of the multiple antibiotic resistance operon(*marRAB*)^40,41^ (Supplemental Figure 3e). These differences in mutational outcomes relative to evolution protocol were also observed in evolved clinical isolate Pb^CEF^ and Pb^CEF-R^ (Supplemental Figure 4f).

### Rationally designed sequential antibiotic treatment results in multidrug survival by resistance, tolerance, and persistence

To design a rational sequence for multiple-antibiotic therapy, we analyzed the outcomes of single-antibiotic treatments. MG^LEV^ cultures remained sensitive to amikacin but had elevated resistance to cefepime, leaving amikacin as a viable continuation for this cohort (Figure 2b, d). Meanwhile, MG^AMI^ cultures had small but statistically significant collateral effects on sensitivity to levofloxacin and cefepime, leaving both antibiotics as viable options (Figure 2c-d). Previous studies have reported on the collateral effects of aminoglycoside resistance toward beta lactam susceptibility^20,25^, so we hypothesized that using amikacin to treat MG^LEV^ cultures would further sensitize them to cefepime as we observed in MG^AMI^ cultures and resensitize them to levofloxacin.

Based on this rationale, we evolved the MG^LEV^ cultures under the selection of amikacin (MG^LEV,AMI^) daily until there was sustained survival, and then we evolved those cultures under the selection of cefepime (MG^LEV,AMI,CEF^) toward the same endpoint (Figure 3a). The evolution protocol, dosing, and durations were the same as their single antibiotic treatment counterparts (Figure 1).

**Figure 3.**
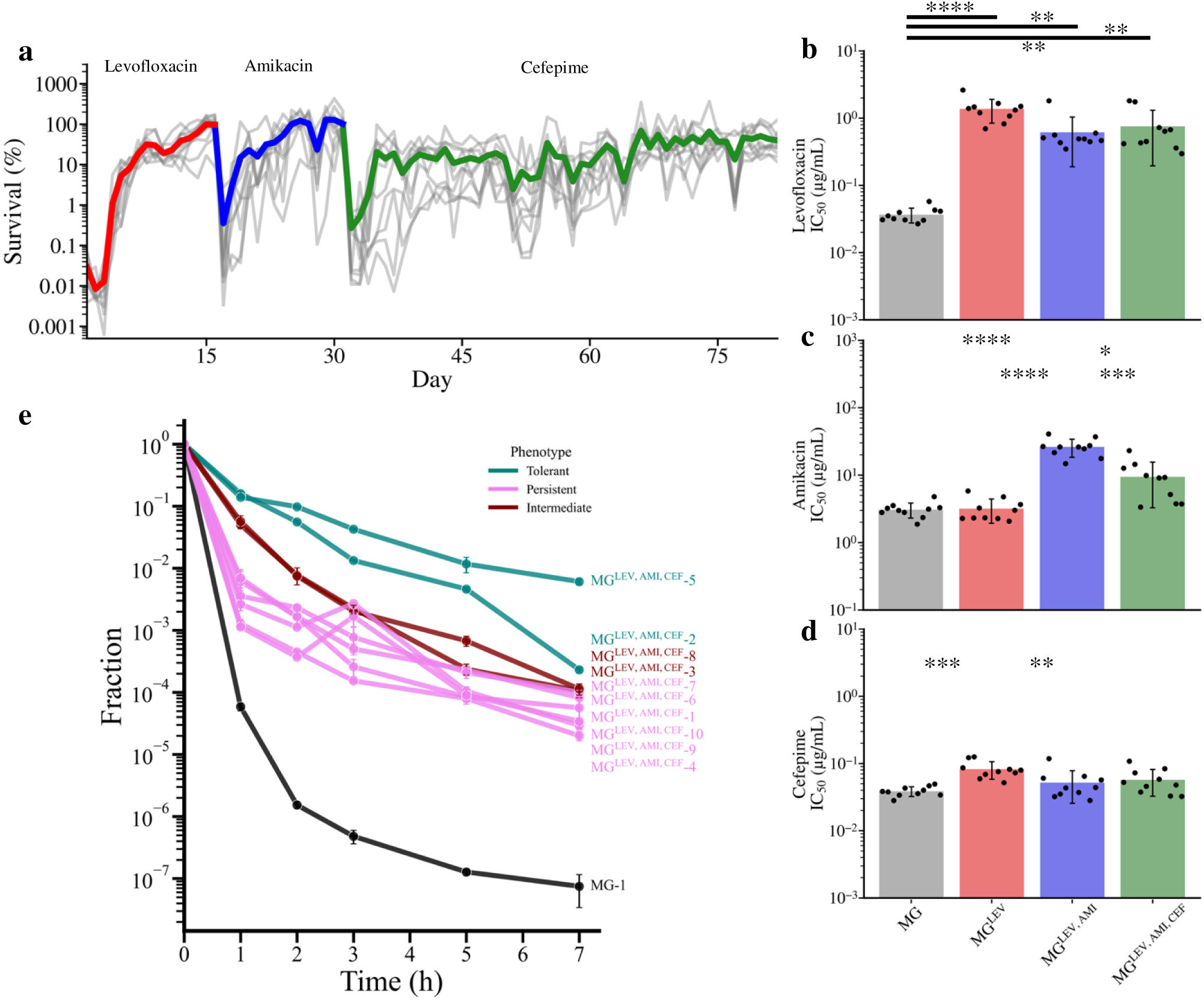
Survival and phenotypic outcomes of sequential antibiotic treatment. a, survival trajectories of bacteria cultures sequentially evolved in the presence of levofloxacin (red), amikacin (blue), and cefepime (green). Gray lines represent individual cultures, and colored lines indicate the mean survival through the respective antibiotic: levofloxacin (red), amikacin (blue), cefepime (green). b-d, bar plots of IC_50_ values in antibiotics levofloxacin, amikacin, and cefepime, respectively, for parental and evolved cohorts (MG: gray; MG^LEV^: red; MG^LEV,AMI^: blue; MG^LEV,AMI,CEF^: green). Bar height corresponds to the mean IC_50_, and error bars reflect the standard deviation. Individual IC_50_ values for replicate cultures re shown as dots. e, MDK assay for sequentially evolved cultures (MG^LEV,AMI,CEF^) showing the median survivor fraction plotted over time, with error bars representing the median absolute deviation (MAD). Line colors differentiate the parental strain (MG: black) and evolved cultures that are characteristically tolerant (teal), persistent (pink), or of a mixed phenotype (maroon).

As anticipated, amikacin caused strong initial killing, with average survival on Day 1 (the 17th cumulative treatment day) of 0.356 ± 0.858% (Figure 3a). However, variability in killing across replicates increased compared to Day 1 of the preceding levofloxacin treatment. This was justified as the terminal MG^LEV^ cultures entered the new treatment with greater genetic diversity than the isogenic parent strain (MG) in our initial experiments (MG to MG^LEV^). The survival trajectories increased rapidly in a sigmoidal pattern (Figure 3a), leading to robust survival. After 15 days of amikacin evolution (31 days in total), the MG^LEV,AMI^ cultures had an average survival of 102.25 ± 56.44 %, so we initiated the cefepime treatment.

The strong killing effect was restored by cefepime on Day 1 (32^nd^ cumulative treatment day) with an average survival of 0.271 ± 0.292 %, and the variability of the killing effect was high (Figure 3a). Like its single antibiotic treatment counterpart (Figure 1f), the survival trajectories through cefepime were gradual and toward an average modest survival of 39.77 ± 34.73 % even after 50 cefepime treatment days (81 days in total).

Evolved bacterial cultures from the final days of each segment of the sequential antibiotic evolution were evaluated for resistance to all three antibiotics (Figure 3b-d, Supplemental Figure 7). Following the amikacin segment of the sequential evolution, MG^LEV,AMI^ cultures became significantly more resistant to amikacin, an ∼8-fold change (p= 4 x 10^-6^) relative to MG^LEV^ cultures (Figure 3c). MG^LEV,AMI^ cultures were also moderately resensitized to levofloxacin relative to MG^LEV^ cultures (∼2-fold, p = 5 ×10^-3^), but MG^LEV,AMI^ cultures remained significantly more levofloxacin resistant than parent MG cultures (p= 2 x 10^-3^, Figure 3b). As we hypothesized, evolution under the selection of amikacin (MG^LEV^ to MG^LEV,AMI^) reversed the cross resistance to cefepime observed in MG^LEV^, resulting in a reduction in the average IC_50_ from 0.082 ± 0.024 µg/mL in MG^LEV^ to 0.052 ± 0.026 µg/ml in MG^LEV,AMI^ (p=8×10^-3^). This IC_50_ value was not significantly higher than the cefepime IC_50_ in the parent MG cultures (Figure 3d).

After evolution under the selection of cefepime (MG^LEV,AMI^ to MG^LEV,AMI,CEF^), levofloxacin resistance did not change significantly relative to MG^LEV,AMI^ (Figure 3b, p = 0.3). However, to our surprise, MG^LEV,AMI,CEF^ was significantly resensitized to amikacin relative to MG^LEV,AMI^ where the average amikacin IC_50_ dropped to 9.42 ± 6.15 µg/ml from 26.23 ± 7.86 µg/ml (p = 10^-4^) (Figure 3c). This phenotypic change was not observed in the single antibiotic treatment counterpart (MG^CEF^). Ultimately, MG^LEV,AMI,CEF^ had no change in cefepime resistance relative to MG^LEV,AMI^ (Figure 3d, p = 0.395). Rather, MDK assays with MG^LEV,AMI,CEF^ cultures again revealed that the increased survival was associated with persistence and tolerance phenotypes (Figure 3e) similar to MG^CEF^ populations. Two MG^LEV,AMI,CEF^ cultures (culture 2 and 5) were characteristically tolerant (Figure 3e, teal lines). Two cultures (numbers 3 and 8) had an intermediate phenotype (Figure 3e, maroon lines). The remaining six cultures (numbers 1, 4, 6, 7, 9, and 10) were characteristically persistent (Figure 3e, pink lines) with a biphasic killing curve. The shift occurred at survivor fractions between 10^-2^ and 10^-3^ indicating relatively large persistent subpopulations (Figure 3e). The mix of persistence and tolerance phenotypes was similar to what was observed in MG^CEF^ cultures (Figure 2e) and Pb^CEF^ cultures (Supplemental 4e).

### Sequential antibiotic treatment drives unique and convergent evolutionary trajectories

Next, we asked how the evolutionary context and history impacted mutational routes toward survival by Illumina whole genome sequencing of frozen samples isolated from evolved cultured at different time points (Figure 4). When we tracked the five GyrA mutations of MG^LEV^ through the subsequent evolutions under the selection of amikacin and cefepime, we observed that GyrA S83L had an advantage in the amikacin treatment regime where it was either fixed or it replaced the previously dominant GyrA mutations (Figure 4a, d, Supplemental Figure 8a). Strikingly, by day 34, S83L was the dominant GyrA mutation across all cultures and all other GyrA mutations went extinct. Mutations of RpoB on the other hand were also selected against during amikacin treatment and were driven to extinction (Supplemental 8b). In all but culture number 2, RpoB mutations became nearly extinct by the conclusion of amikacin treatment (day 31). In culture 2, the *rpoB* indel survived amikacin treatment, but subsequently was selected against through cefepime treatment.

**Figure 4.**
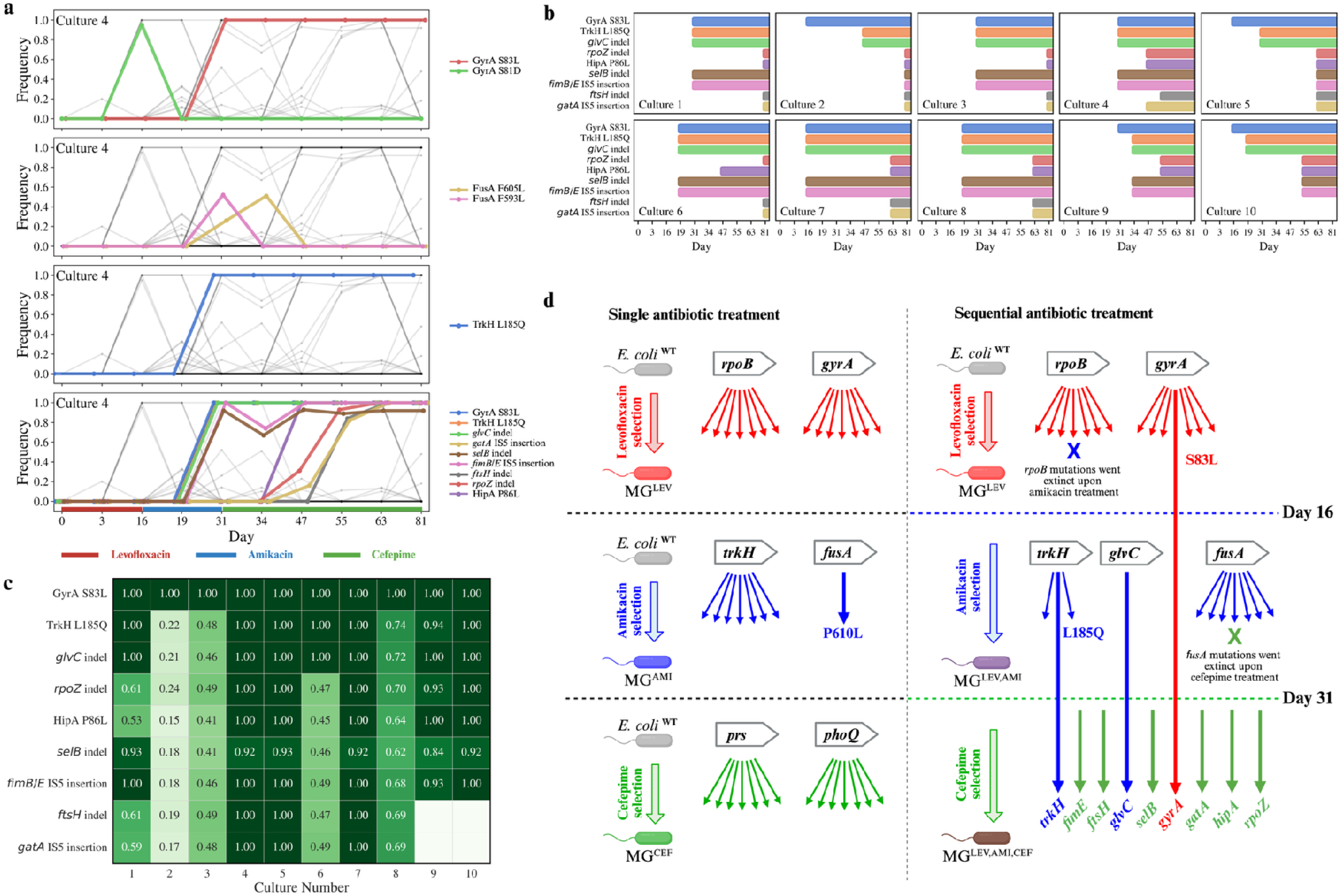
Genotypic outcomes of sequential antibiotic evolution. a, mutation trajectories in Culture 4 of *gyrA* (i), *fusA* (ii), *trkH* (iii) and within the convergent clade mutations (iv). b, emergence and cadence of clade mutations within individual cultures. c, heatmap of final frequencies of convergent clade mutations across replicate cultures. d, cartoon depicting the observed impact of single vs. sequential treatment context on divergent and convergent mutation trajectories (arrows) under the selection of levofloxacin (red) amikacin (blue) and cefepime (green). Arrows are noted with mutations.

Like MG^AMI^, MG^LEV,AMI^ cultures acquired mutations in *fusA* and *trkH* genes, yet the number and type of mutations differed. Where MG^AMI^ cultures had only the FusA P610L mutation, MG^LEV,AMI^ cultures carried 11 unique FusA mutations (including P610L). This difference was most likely due to greater initial population diversity of MG^LEV^ cultures before amikacin selection started. Interestingly, all FusA mutations failed to maintain high frequencies and were subsequently driven to extinction by the final cefepime treatment day (Figure 4a, Figure 4d, Supplemental Figure 8c), indicating a potential fitness cost for FusA mutations under cefepime treatment.

In the case of *trkH*, MG^AMI^ cultures had eight different TrkH mutations, while MG^LEV,AMI^ cultures had only three: T20I, L80Q, and L185Q. MG^LEV,AMI^ cultures possessed high frequency TrkH L185Q mutations which was the dominant allele, a stark contrast to just one MG^AMI^ culture having the TrkH L185Q mutation at a low frequency (0.11). TrkH L185Q quickly fixed within MG^LEV,AMI^ cultures throughout amikacin treatment and maintained in the subsequent cefepime treatment (Figure 4a, Supplemental Figure 8d).

The evolution of survival under the selection of cefepime in MG^LEV,AMI,CEF^ cultures was markedly convergent. In contrast to the genetically divergent evolution observed in MG^CEF^, each of the ten MG^LEV,AMI,CEF^ cultures acquired nearly identical mutations among a set of nine genes at various times (Figure 4b-d, Supplemental Figure 8e). Among the nine mutations were GyrA S83L and TrkH L185Q, which were inherited from previous selection regimens under levofloxacin and amikacin and survived extinction events during subsequent treatment regime changes to amikacin and cefepime, respectively (Figure 4d). The additional mutations emerged with a variable cadence across individual cultures (Figure 4b). Concurrent to TrkH L185Q, a single nucleotide deletion in *glvC*, a poorly characterized putative phosphoenolpyruvate (PEP)-dependent carbohydrate transport system (PTS) component^42,43^, evolved with nearly identical frequencies indicating a common clade and positive selection for the mutated *glvC* (Figure 4d). The deletion occurred at nucleotide 1042 of 1072 generating a downstream stop codon that likely truncates the protein by 17 amino acids.

Next emerged two other concurrent mutations in *selB* and intergenically between *fimB* and *fimE* (fimB/E). *selB* (Figure 4b-d), encoding a translation factor involved in incorporation of selenocysteine into proteins^44^, had a 2-nucleotide insertion that created a frame shift at amino acid 494 and an early stop codon at 526, possibly truncating what is typically a 614 amino acid protein. *fimE*, a regulatory gene that, with *fimB,* controls the production of type 1 fimbriae^45^, was likely lost following an IS5 mobile insertion element inserting between the *fimE* promoter and its transcription start site^46^. This mutation pair was observed in all cultures beside number 2, and either emerged with TrkH L185Q and the mutated *glvC* or after.

All MG^LEV,AMI,CEF^ cultures subsequently had a 7-nucleotide long deletion in *rpoZ*, which encodes the omega subunit of RNA polymerase (Figure 4b-d). The deletion occurs relatively early in the gene (68-74 of 276 nucleotides) and introduces an early stop codon after just 29 amino acids, a severe truncation of what is typically a 91 amino acid protein. In all cultures except culture 2, a mutation was observed in *hipA* (HipA P86L) (Figure 4b-d), a gene named for its association with a <u>hi</u>ghly <u>p</u>ersistent phenotype^47^. Additionally, mutations of *ftsH*, a gene encoding a membrane-bound protease involved in stress response^48,49^, and of *gatA*, a known galactitol-specific component of the PTS system^50,51^ were observed in all cultures besides numbers 2, 9, and 10. The *ftsH* gene incurs an in-phase deletion of 12 nucleotides removing codons 261-264 of 644 within its AAA+ domain. *gatA*, meanwhile is disrupted by the mobile insertion element IS5 following its 69^th^ amino acid (Figure 4b-d). By the final day, many of the discussed mutations had reached high frequencies, and those not fixed shared similar frequencies with others, indicating their concurrence within a common clade (Figure 4b-c).

The shift from divergent evolution under the selection cefepime as a single treatment in MG^CEF^ toward convergent evolution when cefepime was used a sequential treatment in MG^LEV,AMI,CEF^ was striking and not previously reported to best of our knowledge (Figure 4d). These shifts are illustrated in Figure 4d where bacterial populations evolve under the selection of an antibiotic (color) through various mutational options (arrows) given the single vs sequential context of the evolution. Many mutations within evolved populations are driven to extinction within subsequent antibiotic exposures (Supplemental Figure 9).

### Individual convergent mutations are insufficient to produce the observed evolved population level persistence

To evaluate the impact of the commonly observed mutations in MG^LEV,AMI,CEF^ cultures, we introduced each mutation into the wild-type, parental MG1655 (MG) genome (Methods)^52^. The single mutation strains produced from this mutagenesis are referred to going forward as their parent strain background (MG) with the mutated gene in superscript (e.g., MG^gatA^, MG^glvC^).

To evaluate the phenotypic changes in the mutant strains, we measured changes in their exponential growth rate, susceptibility to antibiotics (IC_50_, MIC), and survival by persistence or tolerance in the presence of cefepime (MDK assay, Supplementary Figure 2). The parent strain (MG) had a mean doubling time of 20.07 ± 0.80 minutes (Figure 5a). Mutant strains had limited growth defects. The greatest effects were observed for MG^gatA^ (21.17 ± 0.92 minutes, p = 0.0501), MG^selB^ (21.62 ± 0.41 minutes, p = 0.0017), MG^rpoZ^ (21.91 ± 0.88 minutes, p = 0.0036), and MG^fimE^ (21.66 ± 0.57, p = 0.0026).

**Figure 5.**
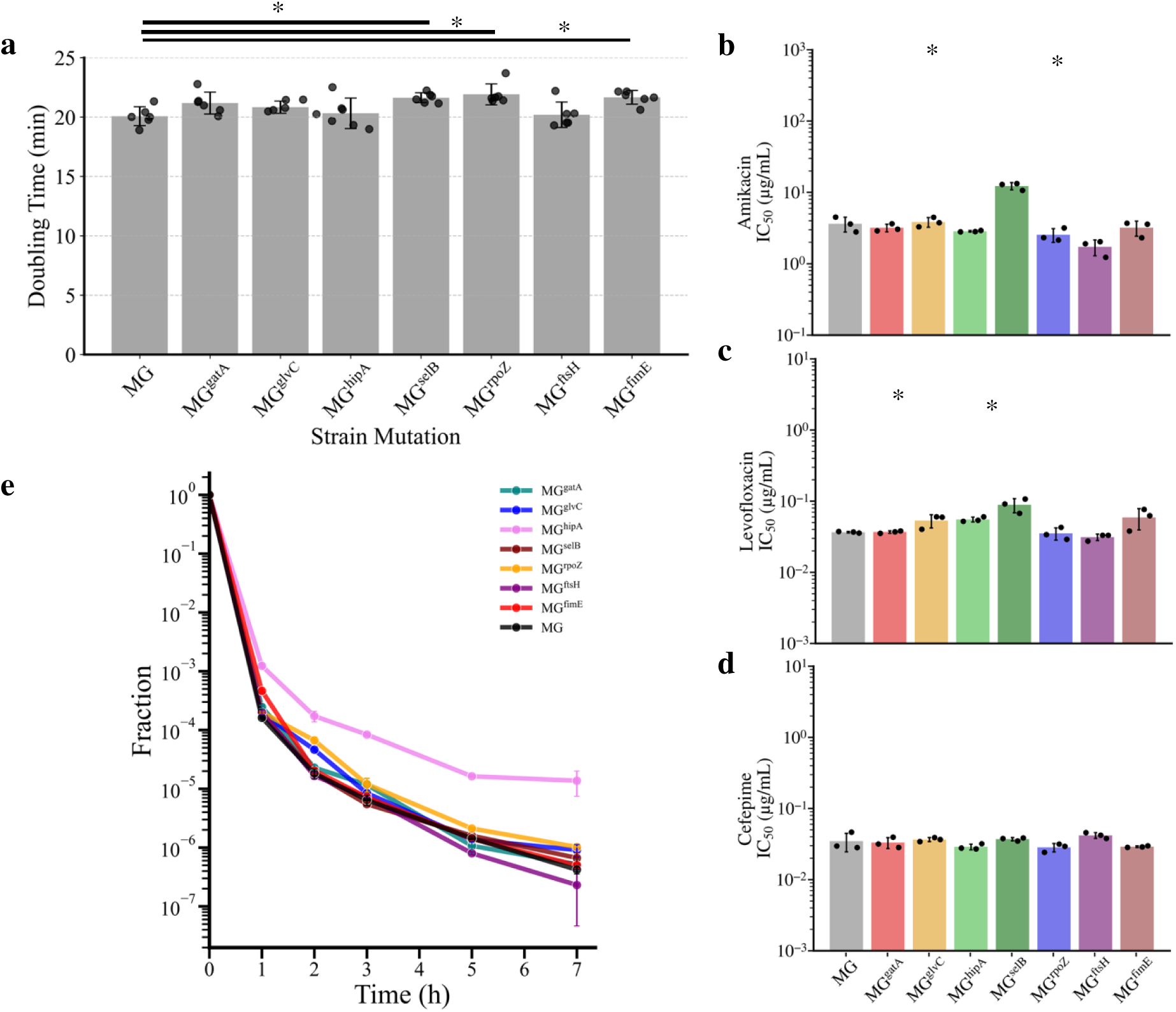
Phenotypic effects of individual convergent clade mutations. a, barplot of doubling times of mutant and parent (MG) strains show mean values of six replicates (dots) with error bar indicating the standard deviation. b-c, IC_50_ of individual mutants in each antibiotic, where bar height shows the mean value for three replicates (dots) and error bar indicates standard deviation. d, MDK of mutants with median survival plotted for three replicates and error bar is the median absolute deviation.

We tested resistance of the individual mutant strains to levofloxacin, amikacin, and cefepime. As before, IC_50_ and MIC were estimated from growth data following overnight growth at 37°C in a range of antibiotic concentrations. Mutants MG^gatA^, MG^glvC^, MG^rpoZ^, and MG^fimE^ did not have any resistance change to any of the three antibiotics (Figure 5b-d). Modest levofloxacin resistance was observed for MG^hipA^ (p=0.014) and MG^selB^ (p=0.046) (Figure 5c). MG^selB^ also had slightly elevated resistance to amikacin (p=0.002). MG^ftsH^ was mildly sensitized to amikacin (p=0.04) (Figure 5b). No changes in cefepime resistance were observed for any of the mutants we engineered.

Next, we used the MDK assay to identify the impact of the mutations on survival by tolerance and persistence. The parent strain (MG) is quickly killed by the effects of cefepime in the first hour with a surviving fraction just above 10^-4^ (Figure 5e). Many of the mutant strains were killed at the same rate as parent MG through each timepoint. Only MG^hipA^ was less susceptible through 1 hour with a surviving fraction of 10^-3^. The rate of killing slowed considerably for MG^hipA^ relative to MG and the other mutants and then continued to have roughly 10-fold greater survival at every time point. Ultimately after 7 hours, MG^hipA^ had a surviving fraction above 10^-5^, while the other cultures were at or below 10^-6^. The biphasic shift in its killing is indicative of persistence, in agreement with previous observations of the phenotype in *hipA* mutants.

The relatively high frequencies and penetrance of the other mutations among the replicate MG^LEV,AMI,CEF^ populations suggest a positive selection in the presence of cefepime. This observation strongly indicates that their contribution to survival and fitness is either epistatic in nature or acts in support of the HipA mutation-led phenotype. Indeed, when the survival of MG^hipA^ is directly compared to that of a diverse, evolved culture, MG^LEV,AMI,CEF^-7, the presence of additional mutations is likely to be associated with enhanced survival. A similar pattern was observed in MG^prs^, a strain engineered with the Prs A114V mutation, and its corresponding diverse, evolved culture, MG^CEF-5^ (Supplemental Figure 10d).

In summary, *hipA* and *prs* mutations display persistence and tolerance phenotypes, respectively. However, evolved cultures carrying the exact same mutations exhibit stronger persistence or tolerance phenotypes, suggesting the presence of epistatic interactions between *hipA* and *prs* mutations and the other mutations observed in the evolved populations.

### Single cell RNA sequencing of evolved cultures reveals metabolic subpopulations and hipA overexpression

To determine how the diverse, accumulated mutations influence bacterial community composition, we used single-cell RNA sequencing (scRNAseq) to characterize changes in the gene expression landscape resulting from sequential antibiotic exposure. Samples were collected from exponentially grown cultures at the same optical density (OD_600_ = 1.0) of the parent strain and two cultures that accumulated largely overlapping mutations: MG^LEV,AMI,CEF^ -4 and MG^LEV,AMI,CEF^-7. Using ProBac-seq^53^, a probe based bacterial scRNAseq method, we analyzed a total of 48,883 single cells from duplicate samples of these three cultures (Supplementary Table 3).

Using dimensionality reduction and graph-based clustering, cells were grouped into 9 unique clusters (Figure 6a). While most cell clusters included at least a few cells from each culture, there were notable instances in which specific clusters were predominantly composed of cells from distinct cultures (Figure 6b, c). The majority of cells in clusters 1 and 2 were of parental origin (>67% and >70% parent cells in each cluster, respectively). Gene-set enrichment analysis (GSEA) identifies translation-related processes and expression of ribosomal subunits as highly enriched in the parental-dominated clusters (Supplementary Table 3, Supplementary Figure 11) These same gene sets were enriched when comparing the wildtype to either culture without cluster-based analysis (Figure 6d, e and Supplementary Table 4). The signature of lower translational activity in the evolved populations is consistent with the common observation that antibiotic persistence is associated with lower activity levels^54^. For example, expression of the 50S ribosomal subunit L10 (*rplJ*) is higher in parent cultures, and significantly upregulated in clusters 1 and 2 (Figure 6 d-f, Supplementary Tables).

**Figure 6.**
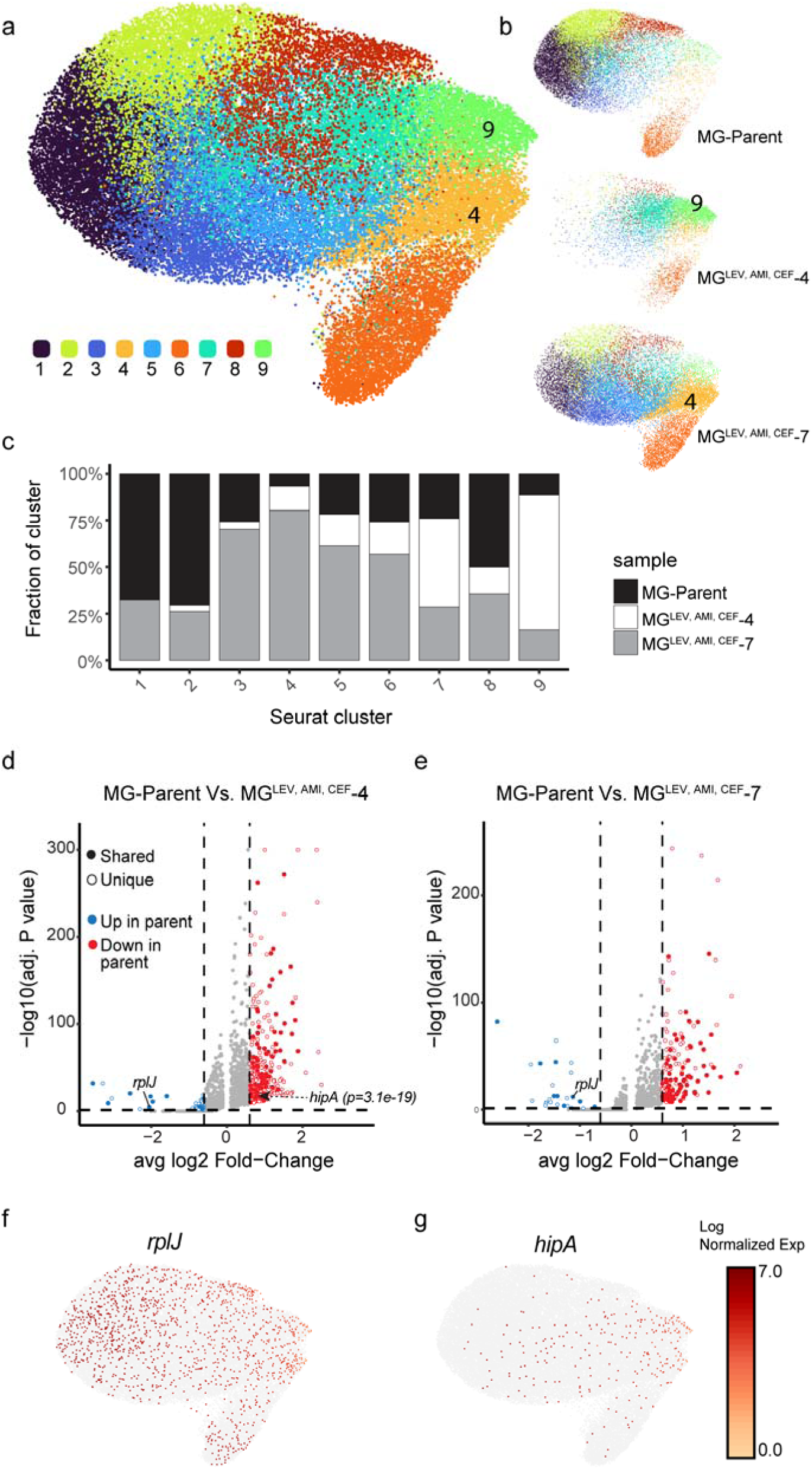
a, UMAP two-dimensional representation of the 9 cell clusters identified in the combined dataset of 48,883 cells. b, UMAP showing cells in either the parental culture (top) or evolved cohorts (middle and bottom). c, Stacked bar chart showing the percent community composition of each cell cluster. d,e, Volcano plot of genes expressed in the parent or MG^Lev,Ami,Cef^-4 (d) and MG^Lev,Ami,Cef^-7 (e). Genes shared between the two evolved strains are denoted by solid markers. Adjusted P values correspond to two-sided Wilcoxon rank-sum test with Bonferroni correction. **f**. Distribution of the 50S ribosomal subunit *rplJ*. **g**. *hipA* is distributed away from clusters 1 and 2 and predominantly within clusters 9 and 4.

Cells from each of the evolved cultures were commonly and significantly enriched in transcripts for the superpathway of glycolysis, pyruvate dehydrogenase, TCA, and glyoxylate bypass, with several glyoxylate shunt genes overexpressed in these cultures (shared genes in Figure 6d, e and Supplementary Table 4). Culture MG^LEV,AMI,CEF^-4 comprised the majority of cells in cluster 9 (>72%), which itself was enriched in this superpathway (Figure 6c, Supplementary Figure 12, Supplementary Table 4). Similarly, culture MG^LEV,AMI,CEF^-7 comprised over 80% of cluster 4 cells, which were enriched in this superpathway (Figure 6c, Supplementary Figure 12, Supplementary Table 4).

In summary, single cell analysis demonstrates that the evolved cultures have cells occupying distinct metabolic niches and contain fewer cells in states that are consistent with high translation activity. This pattern is in line with changes that would be expected from mutations that affect toxin-antitoxin genes including *hipA* identified here. Expression of *hipA* itself was significantly amplified in MG^LEV,AMI,CEF^-4, while in MG^LEV,AMI,CEF^-7 too few individual cells were profiled as having significant overexpression of *hipA* (Figure 6 d-e). The distribution of *hipA* expression shifted strongly toward clusters 4 and 9, key clusters within the evolved lineages (Figure 6g). The observation mirrors the subpopulation survival effect which is characteristic of persistence.

## DISCUSSION

Predicting how bacteria survive antibiotic treatments is crucial for designing ‘evolution-proof’ antibiotics and treatment strategies. Laboratory evolution experiments, whether conducted using *in vitro* or *in vivo* models, have been instrumental in advancing our understanding of antibiotic resistance evolution^55,56^. In this study, we demonstrate that treatment protocol (i.e. high antibiotic peak doses observed in human patients) and context (i.e. whether an antibiotic was administered as a single exposure or in sequence), had consequential effects on phenotypic bacterial survival strategies and the mutational routes toward bacterial resistance.

With high-dose, clinically relevant treatments, we demonstrated that use of levofloxacin and amikacin led to canonical resistance evolution while use of cefepime led to persistence and tolerance evolution, and these phenotypic outcomes were independent of the single or sequential context of the treatments (Figures 2, 3). Likewise, the outcomes of persistence and tolerance were independent of the population structure (isogenic or polyclonal) and bacterial strain (lab strain or clinical isolate) (Figure 2, 3 and Supplemental Figure 4).

We also observed context-specific evolution of collateral antibiotic sensitivity^5,25,26^. Amikacin as a single treatment had only minor collateral effects on sensitivity toward levofloxacin and cefepime (Figure 2c-d), but amikacin as a sequential treatment following levofloxacin substantially reversed the levofloxacin-associated cross resistance toward cefepime (Figure 3b,d). A context-specific evolutionary outcome was also observed with cefepime treatment where cefepime as a single treatment (MG^CEF^) had no effect on cross-resistance to amikacin (Figure 2b), but when used as a subsequent treatment following amikacin, cefepime reduced amikacin resistance of evolved populations (Figure 3c) by selecting against FusA mutations (Figure 4a-ii, Supplementary Figure 8c).

We observed that sequential antibiotic treatment can shift evolutionary trajectories from genetically divergent to convergent. In such scenarios, evolutionary constraints such as epistasis between mutations or fitness costs of mutations in the presence of different antibiotics may provide bacterial populations with fewer viable mutational routes toward resistance in subsequent treatments creating an opportunity for rational treatment design.

When we tested the individual convergent mutations selected under sequential antibiotic treatment for their contributions to survival, we observed that only the HipA mutation resulted in persistent phenotype (Figure 5e). HipA is a kinase that activates persistence and the stringent response by driving the accumulation of the alarmone ppGpp. Further, HipA dimerizes and forms a complex with a HipB dimer to repress the *hipBA* operon, and mutations have been shown to weaken the interaction, allowing greater free HipA and less transcriptional autoregulation of the operon^57–61^. While MG^hipA^ was the only mutant strain with increased survival by persistence, the degree of its survival throughout the MDK assay was lower relative to the survival observed in MDK assays of evolved MG^LEV,AMI,CEF^ (Supplementary Figure 10). We hypothesize that the additional convergent mutants contribute to survival through epistatic interactions among them. Among them, mutations in *ftsH* and *rpoZ* may have the clearest connection to the persistent state associated with the HipA mutant.

FtsH was previously shown to degrade stress response regulators to allow bacteria to return to a prestressed state^48,49^, and degrade the LpxC and KdtA enzymes that are involved in the early steps of LPS biosynthesis^62^. Interestingly, the FtsH-mediated degradation of LpxC is modulated by ppGpp^63^. The 4 codon deletion within the AAA+ domain of FtsH observed here may alter ATP binding or protease efficiency as a way of adapting to prolonged stress or to compensate or potentiate the HipA persistence phenotype^64^.

RpoZ, the non-essential omega subunit of RNA polymerase, interacts with other RNA polymerase subunits which creates a binding interface for ppGpp that contributes to the propagation of the stringent response and persistence phenotype^65,66^. The proposed connection to HipA is further supported in the observed cadence of mutation emergence, where mutations of FtsH and RpoZ always coincide or succeed the mutation of HipA throughout evolution under sequential antibiotic treatment.

The results of our single cell RNA sequencing analysis reveal information about transcriptional complexity within the evolved populations. Clustered cells with widespread expression changes of metabolic and translation programs highlight additional opportunities to characterize bacterial susceptibility following distinct treatment regimes. It remains unclear the exact mechanistic role for the transcriptional shifts in superpathway genes, but it may likely lead to metabolic shifts affecting growth rates and cell states which contribute to antibiotic survival. Our analysis revealed reduced expression of ribosomal genes and overexpression of *hipA* in subpopulations of evolved cells consistent with the biphasic killing effect observed in MDK assays, despite the high frequencies of the mutation. (Supplemental Figure 10d). Akin to the allele-specific susceptibility and survival we observed, we hypothesize that some bacterial subpopulations differentially expressing distinct gene sets will have greater survival during antibiotic treatment.

This work offers an initial roadmap for identifying sequential antibiotic treatments that promote favorable mutational trajectories in the face of multiple treatment failures. Our ability to predict mutational routes toward resistance, persistence, and tolerance may improve antibiotic choice based on known collateral phenotypic effects. Further, as shown here, sequential treatments can shift evolution from divergent to convergent routes where bacteria may have fewer mutational options to surviving subsequent treatments, marked by extinction events and variable allelic diversity. In pursuit of rational treatment design and improved clinically efficacy, future investigations of this kind could be performed experimentally, as was done here, with more antibiotics and sequences, or rather by sequencing clinical isolates following sequential antibiotic treatment courses.

## Materials and Methods

### Bacterial strains

Lab strain *Escherichia coli* (MG1655), a genomically barcoded, clinical isolate *Escherichia coli* (PbEc)^56^, and mutant MG1655 clones generated by oligo recombineering were routinely cultured in lysogeny broth (LB) medium (RPI) aerobically (shaking at 230 rpm) at 37°C. When applicable, growth was determined by spectrometry (OD_600_), and survival was determined by enumeration of cultured colony forming units (CFU) on LB agar plates (RPI). Plate-based assays (MIC) were performed in LB medium with aerobic growth (shaking at 400 rpm) at 37°C and controlled humidity (80%).

### Antibiotic Stock Preparation

Concentrated stocks of antibiotics levofloxacin (Alfa Aesar) and cefepime (Apotex) were prepared in water, filter sterilized, and stored at -20°C for single use, each 10mg/mL. Amikacin sulfate (Heritage) was diluted as needed from purchased liquid stock.

### Colony forming unit enumeration

Multiple assays relied on the enumeration of colony forming units. To perform these measurements, each well of 96 well plates was filled with 180 µL of phosphate buffer saline (PBS, Sigma) solution. 20 µL samples of cultures were then serially diluted (1:10) across the rows of 96 well plate up to a 10^-6^ dilution. 20 µL of each dilution was plated on LB agar plates in lines, and agar plates were incubated overnight (37°C). The following day, individually colonies were counted from the plated dilutions with the maximum countable colonies. When necessary, cultures or aliquots were centrifuged and washed with PBS to remove antibiotic prior to diluting and plating.

### Evolution Assays

The evolution assay developed presently was repeated daily beginning with a 1:10 dilution of overnight cultures into 10 mL of antibiotic-free LB media. Diluted cultures were grown for 1 hour. Prior to treatment, a 20 µL sample was added to the 96 well plates prepared for CFU enumeration. The remaining culture was dosed with antibiotic and returned to shaking incubation. Diluting and plating related to CFU enumeration was performed during the incubation period. Incubation times were previously optimized to achieve 99.9% killing on the initial day. After the necessary incubation time (1 hour for levofloxacin and amikacin; 2 hours for cefepime), bacteria were pelleted by centrifugation (6000g, 5 minutes) and washed twice with PBS to remove the antibiotic. The cleaned pellets were resuspended with the equivalent volume of antibiotic-free LB media, and a post-treatment 20µL sample of the culture was collected for CFU enumeration. Cultures were promptly returned to shaking and incubation for overnight growth. The protocol was repeated daily until an increased sustained survival was observed. Percent survival was calculated by dividing post-treatment CFU counts by pre-treatment CFU counts. Sequential antibiotic treatments immediately followed the previous antibiotic treatment.

For the conventional laboratory evolution protocol, overnight cultures were diluted in antibiotic-free LB media (1:100). Antibiotic was serially diluted (1:2) across 5 mL LB media in 15 mL conical tubes to prepare a range of concentrations surrounding a predetermined MIC (MIC/4, MIC/2, MIC, 2xMIC, 4xMIC). Parental strain bacteria from the overnight growth were added (1:500) to tubes containing antibiotic and then grown aerobically overnight as described previously. The following day, the tube with the highest antibiotic concentration that allowed for robust growth was used to inoculate (1:500) freshly prepared conical tubes with an increased range of antibiotic concentrations relative to the prior day, centered around the passaged concentration (e.g., MIC/2, MIC, 2xMIC, 4xMIC, 8xMIC). This was repeated daily.

Routinely, bacteria were collected and saved as a glycerol stock by combining 500 µL of bacteria culture with 500 µL 50% glycerol.

### Resistance Determination

In 96 well plates, twofold dilutions series of antibiotics were prepared in 100 µL of LB media. Then, 100 µL of diluted bacteria cultures (OD_600_: 0.01) was added. Each plate included “Media Only” wells to measure background OD signal and “Cells Only” wells to assess maximum growth in the absence of antibiotic. Plates were incubated overnight at 37°C, shaking at 400 rpm, in a humidified (80%) incubator (Infors HT). Following incubation, OD_600_ was measured (Biotek Epoch2 plate reader), and raw OD values were exported and processed using a custom Python script.

First, a global or plate-specific median background was computed from the “Media Only” wells. This median value was subtracted from the raw data to correct for background signal. Next, each strain–antibiotic dataset was normalized by dividing the background-corrected OD values by the median OD of its corresponding “Cells Only” wells, yielding fractional growth that ranged from near 0 (complete inhibition) to 1 (no inhibition). Using these normalized data, a two-parameter logistic (2PL) curve was fit to each strain–antibiotic combination. Specifically, the present function was employed to model growth

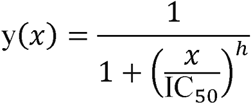

where x is the antibiotic concentration, IC_50_ is the concentration at which growth is half the no-antibiotic control, and h is the Hill slope reflecting the steepness of the inhibition curve. Experiments were conducted with technical triplicates across 12 antibiotic concentrations, ensuring sufficient data points across multiple conditions for robust curve fitting.

Parameter optimization was performed via SciPy’s least_squares routine with a robust “dogbox” algorithm and “soft_l1” loss, which reduces sensitivity to outliers. The minimum inhibitory concentration (MIC) was then defined as the concentration at which growth reached 5% of the no-drug control (i.e., 95% inhibition). This value was obtained by numerically solving the fitted 2PL function at y=0.05. To quantify uncertainty in IC_50_ and MIC estimates, a bootstrap analysis was performed by resampling each dose–response dataset 1,000 times with replacement and re-fitting the curve. The 2.5^th^ and 97.5^th^ percentiles of these bootstrap distributions defined 95% confidence intervals, providing robust estimates of parameter variability. When the model-predicted IC_50_ or MIC exceeded the maximum tested antibiotic concentration, an “insufficient drug” flag was assigned, indicating that the true inhibitory concentration likely lay beyond the tested range, and excluded from further analysis. All original code is available in this paper’s supplemental information.

### Minimum duration of killing (MDK) assay

Overnight *E. coli* cultures were diluted 1:10 in antibiotic-free LB media and grown aerobically for 1 hour. Initial CFUs were measured as described previously. The cultures were then exposed to 129 µg/mL of cefepime for 7 hours. At hours 1, 2, 3, 5, and 7, 500 µL aliquots were collected in 1.5 mL microfuge tubes, centrifuged, and washed twice with PBS to remove the antibiotic. Pellets were resuspended in 500 µL PBS. At hours 5 and 7, final pellets were resuspended in 50 µL PBS, 20 µL of which was plated and 20uL was diluted further, to increase the limits of detection. CFUs were enumerated, and survivor fraction was calculated by dividing the ongoing counts by the initial CFU measurement.

### Bacterial population sequencing

Bacteria cell pellets of antibiotic-evolved, passaged untreated control, and parental cultures were prepared from overnight cultures and sent to Seqcenter (Pittsburgh, PA) for DNA extraction (ZymoBIOMICS) and whole genome sequencing (Illumina). Samples were sequences using 150 base pair paired-end reads. For evolved cultures, approximately 2 giga base pairs of data per sample was received yielding coverage of least 100x throughout the genome. Sequencing data was processed using breseq (Barrick lab), comparing reads to a MG1655 reference genome ^67^. Alleles present in the passaged, untreated controls were tracked but were removed from the dataset of evolved populations. Likewise, alleles present in the parental strain relative to the reference genome were subtracted. Remaining mutations were filtered to include only those which exceeded 10% at any timepoint and these alleles were traced across timepoints. Commonly mutated genes were presented for each single antibiotic evolution with full mutation sets presented as supplemental figures. The full breadth of mutations within the cohorts of evolved cultures following single and sequential evolutions are shown in Supplemental Figure 9, with the exlusion of MG^LEV,AMI,CEF^ cultures 2, 3, and 8 which had exceedingly numerous mutations associated with a hypermutator mutation of *mutL*. Further, the *mutL* mutation was first identified in culture number 2 during the amikacin regime. Later, on day 81, *mutL* was observed in cultures 3 and 8 along with other mutations present in culture 2, making it difficult to rule out a possible contamination of those cultures sometime between days 63 and 81.

### Oligo recombineering

Chromosomal mutations were made using oligo recombineering, and gene deletions were made using oligo recombineering followed by Bxb-1-Integrase-Targeting (ORBIT). These methods each incorporate mutations encoded on transformed oligonucleotides through homologous recombination during DNA replication, and require elements encoded on a helper plasmid^52^. Parental MG1655 bacteria were made electrocompetent and then transformed with the pHelper_Ec1_V1_gentR (pHelper-V1) for oligo recombineering or with pHelper_noMutL_V2_gentR (pHelper-V2) for ORBIT. Selection was made possible by gentamicin. pHelper-V1 expresses single stranded DNA annealing proteins (SSAP), CspRecT, and a dominant negative MutL, which improves editing efficiency by suppressing mismatch repair^68^. Sucrose counterselection is made possible by *sacB* gene encoded on the plasmid. The expression is driven by the induction of the XylS/m-toluic acid system. pHelper-V2 differs in that it does not include the dominant negative MutL. Transformed bacteria were induced with toluic acid for 1 hour prior to storage at -80°C.

For simple oligo recombineering, 90-nucleotide long oligonucleotides were designed with the desired mutations at the center flanked by homology regions within the gene, and each was ordered from Integrated DNA Technologies (IDT) (Table1). Oligonucleotides were prepared and diluted to 25 µM in water. 50 µL of MG1655-pHelper-V1 was mixed with 2 µL of diluted oligonucleotide, electroporated (1.8mV), and recovered in LB media for 1 hour at 37°C. Recovered bacteria were steaked on LB agar for single colonies and incubated overnight at 37°C.

Individual colonies were picked into 100 µL PBS in 96 well plates for screening by polymerase chain reaction (PCR). Throughout screening, colonies were routinely substreaked on LB-sucrose (7.5%) plates to drive the removal of the pHelper-V1 plasmid. Plasmid loss was confirmed by plating on LB-gentamycin (15 µg/mL) plates.

### Preparation of electrocompetent cells

MG1655 cultures were made electrocompetent prior to transformations with plasmids pHelper-V1 and pHelper-V2, and subsequently MG1655-pHelper-V1 and MG1655-pHelper-V2 were made electrocompetent prior to transformations with oligonucleotides. Each approach varied slightly.

Beginning with plasmid transformations, filtered water was aliquoted in 50 mL conical tubes and refrigerated overnight. MG1655 was cultured overnight in 10 mL LB media. The following day, a 1:100 dilution of the overnight was performed into a fresh 10 mL LB and grown for 2 hours. During this time, the water aliquots, empty 15 mL conical tubes, and microfuge tubes were placed on ice. After 2 hours of growth, the culture was split into two, 15 mL tubes. These cultures were centrifuged at 4000g, 2°C, for 10 minutes. The pellets were washed twice with 10 mL of cold water. The final supernatant was fully removed by pipetting. Pellets were then resuspended in 1 mL of cold water, combined, and then split into 2 microfuge tubes. The microfuge tubes were centrifuged for 10 minutes, and each pellet was resuspended in 100 µL of cold water. In separate tubes, 100 µL of the culture was combined with 20 ng of either plasmid and each was electroporated (2.5 kV). Immediately, 1 mL of prewarmed recovery media (NEB) was added and moved to a 15 mL conical tube where it recovered for 1 hour at 37°C in the shaking incubator. Following its recovery, 200 µL of each culture was plated on LB agar plates containing 15 µg/mL gentamycin for positive selection of transformants. A non-transformed culture was used as a control. After overnight incubation, single colonies were picked and grown overnight in liquid culture with gentamycin (15 µg/mL).

The following day, the cultures, transformed MG1655-pHelper-V1 and MG1655-pHelper-V2 were induced and made competent again. Filtered water and 10% glycerol had been refrigerated overnight. From the overnight cultures, a 1:1000 dilution was made into 200 mL of LB with gentamycin and grown for 3-4 hours to ∼ 0.3 OD. While growing, reagents and tubes were placed in ice. When the cultures reached 0.3 OD, the cultures were induced with 1 mM toluic acid (added from a 1000x stock solution) and incubated for 30 minutes. Following incubation, the cultures were removed and placed in an ice bath with occasional swirling to stop their growth. Then 50 mL aliquots of the cultures were transferred to iced conical tubes. These were centrifuged at 5000g for 10 minutes at 2°C. The supernatant was poured off, and pellets were resuspended with 45 mL cold water. Resuspended pellets were centrifuged, and pellets were resuspended with 10% glycerol. This was repeated twice more, but the final pellets were resuspended in the residual glycerol. Cultures of the same strain were pooled and then aliquoted (100 µL) into small microfuge tubes. These were chilled on dry ice for 5-10 minutes before being stored at - 80°C. These resulting stocks were competent and induced to allow for the transformation and incorporation of targeting oligonucleotides.

Cultures transformed with oligonucleotides were plated on non-selective plates, and thus colonies required screening by PCR and then confirmation by Sanger Sequencing (Eurofins). For each mutant strain prepared, full-length and mutant-specific PCR primers were designed (Table 2). The mutant-specific primers relied on the small differences in sequences and were paired with the full-length primer in the opposite direction. PCR reactions were performed using GoTaq polymerase (Promega), the appropriate primer set, water, and a 1 µL inoculation of the colony-PBS suspension. Prior to screening, the melting temperatures for reactions using mutant specific primers were optimized by gradient PCR of parent strain and a pool of presumptive mutant colonies. Then, batches of individual colonies were screened by mutant-specific PCR. Mutant PCR reaction products were routinely run on 1% agarose gels with ethidium bromide and imaged under ultraviolet light (Syngene G:BOX Chemi XT4 with GeneSys software). Positive colonies were then PCR amplified using full-length primer pairs. These full-length PCR products were cleaned (Nucleospin Gel and PCR cleanup kit) and submitted for Sanger sequencing (Eurofins).

Analysis of the sequencing results often revealed mixed colonies with peaks representing mutant and wild-type nucleotides. When this occurred, positive colonies were substreaked (LB-sucrose), and rescreened. After several iterations, Sanger sequencing results were pure, and the mutational status was confirmed by sequencing further subclones for purity. For the larger mutations made by oligo recombineering, the efficiency was lower and required a pooling approach to minimize reagent use and increase throughput. When mutants were confirmed and purity achieved, single colonies were picked and grown in LB overnight and then saved as stocks in 25% glycerol.

### Bacterial growth assay

Strains were cultured overnight from glycerol stocks in 10 mL LB media. The following day, a 96 well plate was prepared with 100 µL LB media in each well. Overnight cultures were diluted in fresh media to an OD_600_ of 0.1. To media-only wells, an additional 100 µL LB media was added. To culture wells, with six replicates of each strain, 100 µL of diluted culture was added. The plate was promptly incubated in Biotek Epoch2 plate reader and grown for 20 hours at 37°C with shaking. Every 2 minutes, OD measurements were made. Growth curves were analyzed using a custom Python script. OD readings were first filtered to values between 0.02 and 0.08 to capture the log growth phase. Linear regressions were performed on the log transformed OD values within this range as a function of time. The resulting slope was converted to a natural log-based growth rate. Doubling time was then calculated using the formula

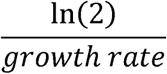

and reported in minutes. The scipy.stats module was used to calculate means, standard deviations, and to perform t-tests of doubling times.

### Cell fixation for single cell RNA sequencing

Overnight cultures of strains were inoculated (100uL in 10mL) from frozen stocks, avoiding population bottlenecking. Cultures were diluted to 1 OD, and 1 mL of culture was fixed with 67µL of paraformaldehyde through a 30 minute incubation. Paraformaldehyde was removed by centrifugation (5 min, 6000 rcf). Pellet was resuspended with 1 mL of 0.2x SSC Buffer (Invitrogen). Buffer was removed by centrifugation, the pellet was resuspended in 350 µL of MAAM (4 parts methanol, 1 part acetic acid), and briefly stored at -20°C.

### Single cell RNA sequencing

Fixed cells were processed using standard ProBac-seq protocols as follows: 100 ul of cells in MAAM solution were centrifuged at 6000 rcf and washed 3 times in 300 ul PBS solution. After PBS washes cells were permeabilized by incubation for 30 minutes with 250 units of lysozyme (ReadyLyse, LGC BioSearch) in 100ul PBS. Cells were centrifuged again at 6000 rcf and washed with 300ul 1x PBS with 0.1% tween solution. Cells were again centrifuged and washed in PBS to remove the detergent and resuspended in Probe Binding Buffer (100□µl of probe binding buffer consisting of 5□×□SSC, 30% formamide, 9□mM citric acid at pH 6.0, 0.1% Tween 20, 50□ug/ml heparin and 10% low molecular weight dextran sulfate). Cells were incubated for 30 minutes at 50°C before 25 ul of probes were added at a concentration of 1,200 ng/ul, and the mixture was left to incubate overnight at 50°C with shaking at 300 RPM. Following incubation cells were washed 7 times in probe-wash solution at 50°C (5□×□SSC, 30% formamide, 9□mM citric acid pH 6.0, 0.1% Tween 20 and 50□ug/ml heparin) and finally washed twice in room temperature PBS.

Single-cell microfluidic encapsulation was done using a 10X Genomics chip-G and a Chromium Controller with the Chromium Single Cell 3′ Reagents Kit (v3 chemistry) as described by 10X Genomics using a protocol modified to achieve bacterial scRNA-seq described previously ^69,70^. For encapsulation a master mix containing the following reagents was prepared: 33□µl of 4X ddPCR Multiplex Supermix (BioRad), 4□µl of in-drop PCR primer (10□µM), 2.4□µl additive A (10X Genomics) and 26.8□µl dH_2_O. Prepared cell samples were diluted to 1,000 cells µl^−1^ and loaded as a total of 16,000 cells per condition to achieve a targeted cell recovery of 10,000 cells. After encapsulation each sample was transferred to an individual PCR tube and cycled 6 times using the following thermocycler program: 94□°C for 5□min (before cycling), 94□°C for 30□s followed by 50□°C for 30 s then 65□°C for 30□s for 6 cycles and a final hold at 12□°C.

Libraries for NGS sequencing were completed using two additional PCR reactions as detailed previously ^69,70^ and sequenced using an Element AVITI 150 cycle kit using paired end sequencing of 8 bp for the i7 index, 28bp for Read 1 and the remaining cycles for Read 2.

Reads were trimmed and the probe-based UMI was upended in place of the traditional 10X-bead provided UMI using cutadapt in order to allow for true molecular identification, as detailed previously ^69,70^. Samples were mapped and aggregated using the 10X cellRanger pipeline. Analysis of single cell libraries was done using the R-based Seurat pipeline ^71^. Standard parameters in Seurat were used for data normalization and to identify the variable features. Louvain clustering was used with resolution set to 0.7 using Seurat FindClusters function. We tested a range of resolutions from 0.5 to 1.5 and find that the main clustering features presented in the text remain across this range of clustering parameters. The Seurat function FindMarkers was used and a log-fold change cutoff of 0.5 chosen to highlight highly differential genes in the main-text volcano plots and a log-fold change cutoff of 0.25 for the supplementary volcano plots in the Supplementary Figures. The full output for all differentially expressed genes is shown in the DGE Supplementary Tables.

### Quantification and Statistical Analysis

Statistical analyses of MIC data were performed using Python (v3.10) with the scipy.stats package. Means and standard deviations of individual cultures within cohorts were calculated. Student’s t-test was used, assuming a normal distribution, to compare the means of relevant groups, treated vs parent and among segments of the sequential antibiotic treatments. Results are reported as mean ± standard deviation with their corresponding p-values. Significance p-value thresholds were noted with asterisks for p<0.05 (*), p<0.01 (**), p<0.001 (***), and p<0.0001(****). Minimum duration of killing (MDK) assay data was reported as median ± median absolute deviation.

## ACKNOWLEDGEMENTS

This study was supported by the Advanced Research Projects Agency for Health (ARPA-H) (1AYSAX000005-01), NIH grant R01GM125748, UTSW High Risk High Impact Award, and Welch Foundation I-2082-20240404.

---

**Supplemental Figure 1.**
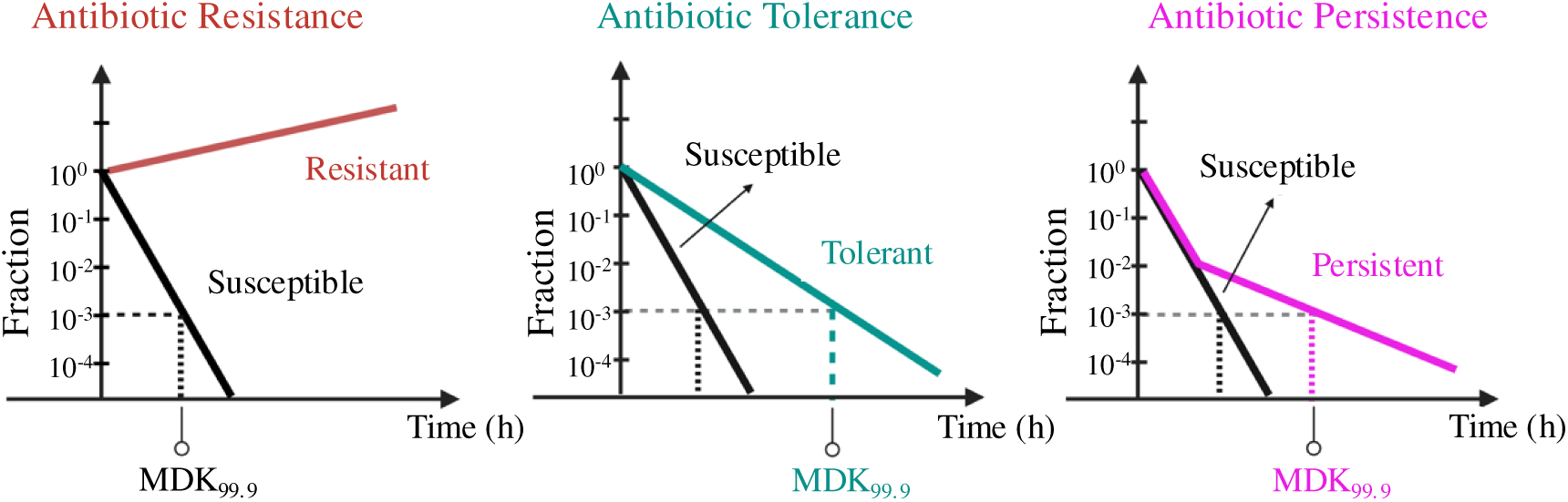
Differences among survival strategies in time kill assay. Illustration of the MDK curves expected for bacteria which are resistant (red), tolerant (teal), persistent (pink), and susceptible (black). Differences include shifts in the minimum duration for killing 99.9% of bacteria (MDK_99.9_) and the biphasic nature of the MDK curve in persistent populations.

**Supplemental Figure 2.**
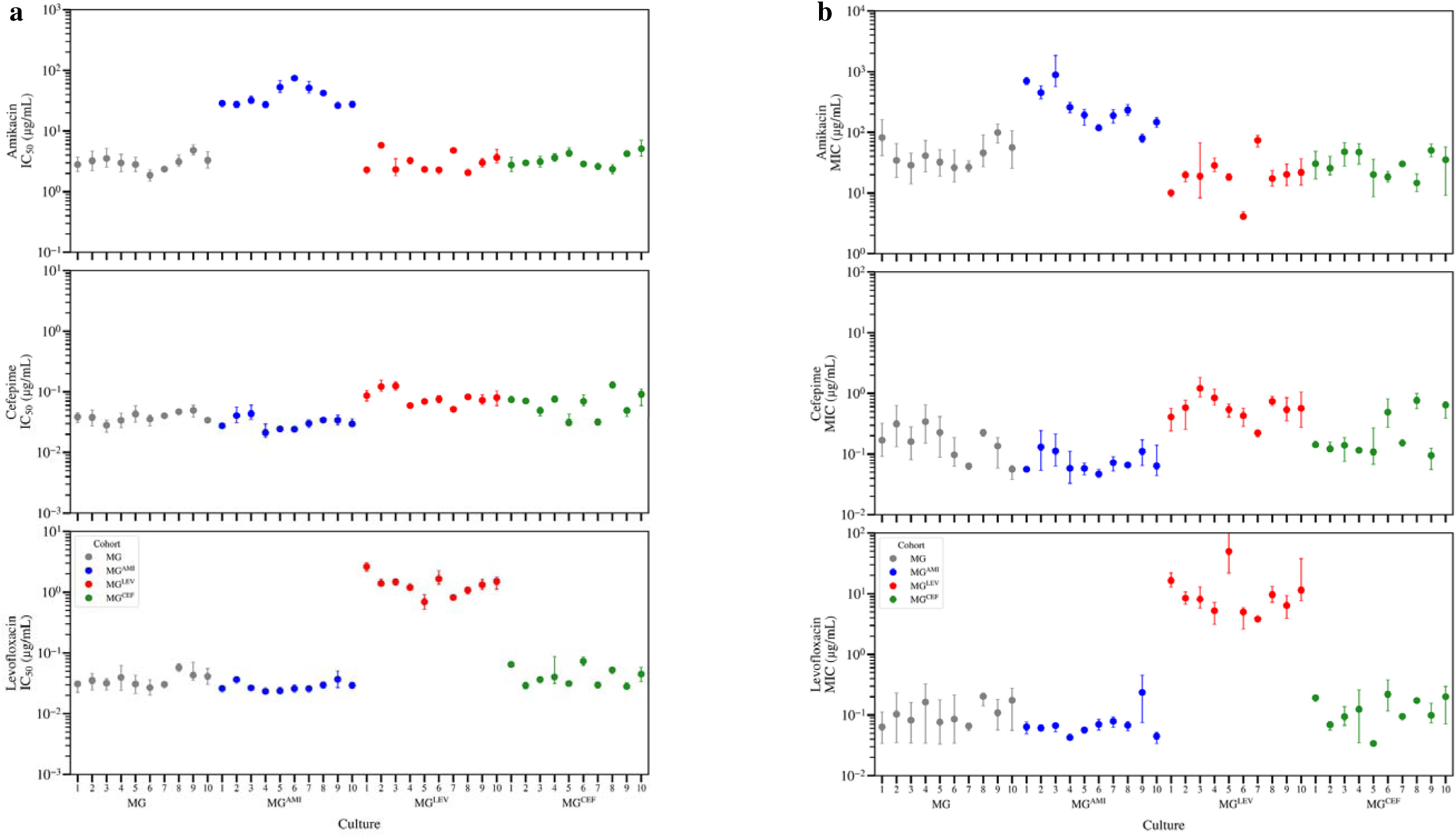
IC50 and MIC values of individual evolved cultures. a, Average IC_50_ values (error bars: standard deviation) for individual cultures of parental strain (MG, gray) and cultures evolved in the presence of amikacin (MG^AMI^: blue), cefepime (MG^CEF^: green), and levofloxacin (MG^LEV^: red). b, Average MIC values (error bars: standard deviation) for individual cultures of parental strain (MG, gray) and cultures evolved in the presence of amikacin (MG^AMI^: blue), cefepime (MG^CEF^: green), and levofloxacin (MG^LEV^: red).

**Supplemental Figure 3.**
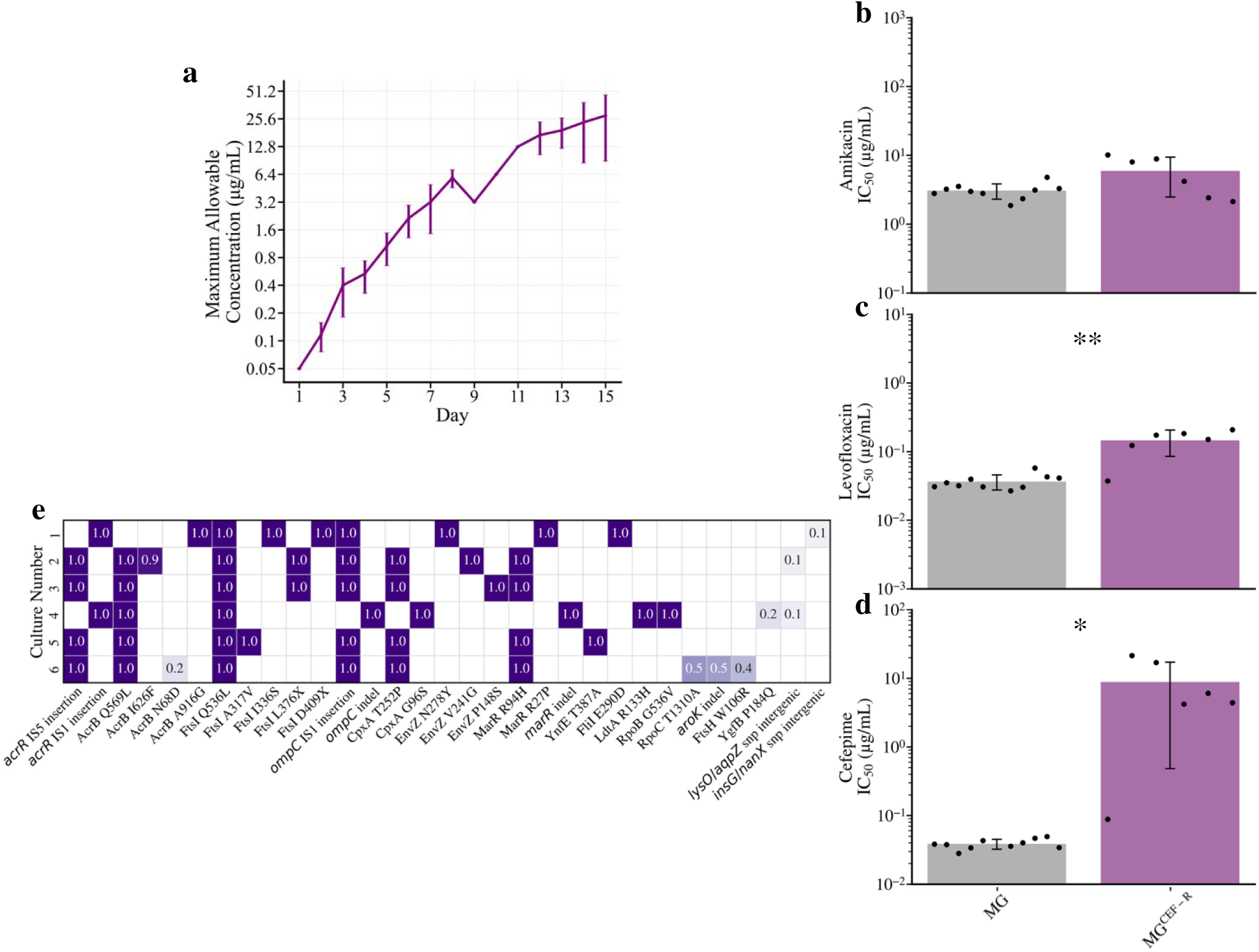
Phenotypic and genotypic outcome of cefepime resistance evolution protocol in lab strain. a, Maximum allowable concertation for growth over time in conventional evolution in the presence of cefepime. Plotted line represents the mean concentration and error bars represent the standard deviation. b-d, bar plots of IC_50_ values in antibiotics amikacin, levofloxacin, and cefepime, respectively for parental and cefepime-resistance evolved cohorts (MG: gray; MG^CEF-R^: purple). Bar height corresponds to the mean IC_50_, and error bars reflect the standard deviation. Individual IC_50_ values for replicate cultures are shown as dots. e, heatmap of selected mutations and their frequencies among individual cultures of MG^CEF-R^.

**Supplemental Figure 4.**
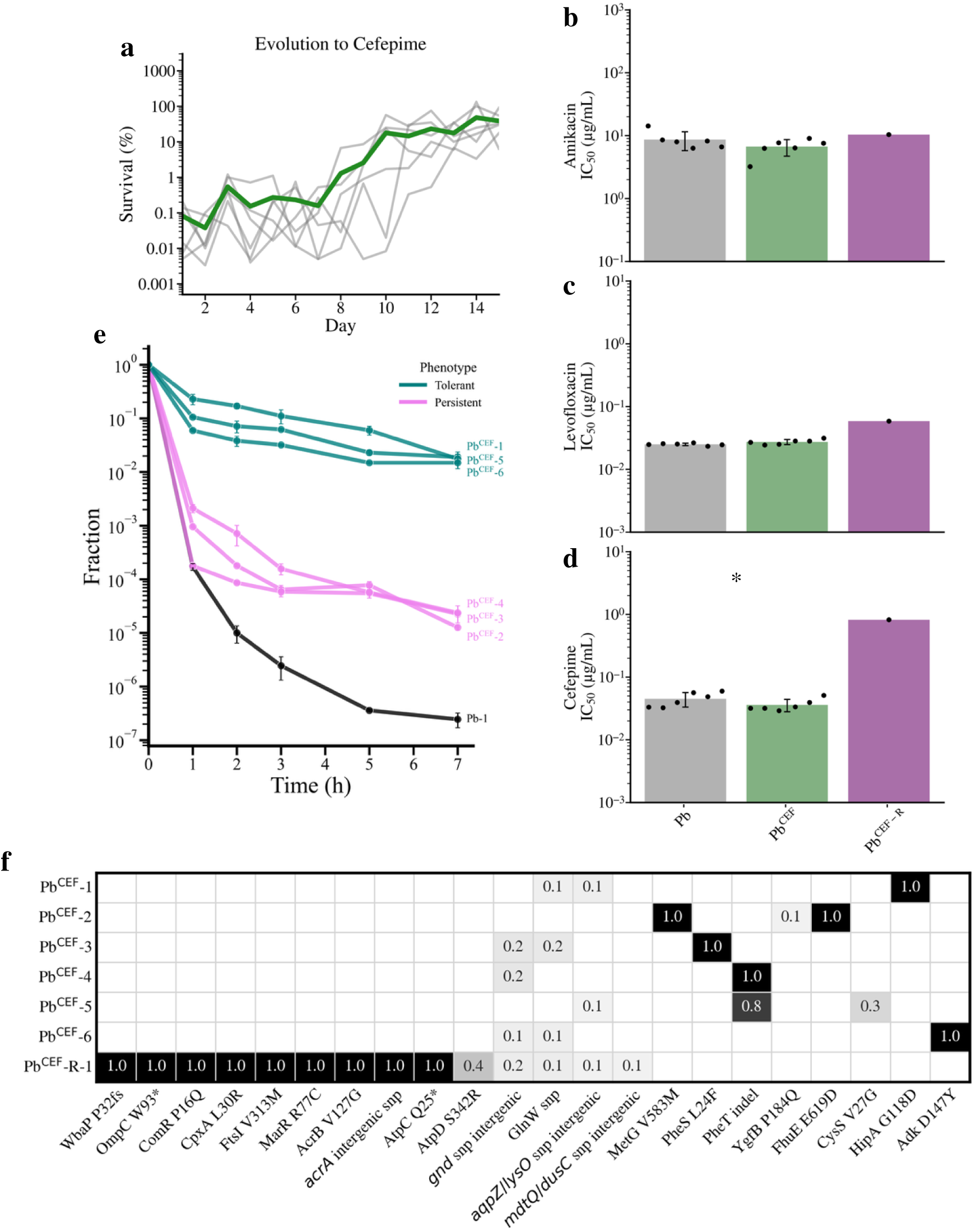
Phenotypic and genotypic outcomes of cefepime evolution in clinical strain. a, Survival trajectories of clinical bacterial cultures evolved in the presence cefepime. Gray lines represent individual replicate cultures, and the mean survival is shown in green. b-d, bar plots of IC_50_ values in antibiotics amikacin, levofloxacin, and cefepime, respectively, for unevolved clinical strain (Pb: gray), cefepime-evolved clinical strain using our evolution protocol (Pb^CEF^: green) and cefepime-evolved clinical strain using the conventional evolution protocol (Pb^CEF-R^). height corresponds to the mean IC_50_, and error bars reflect the standard deviation. Individual IC_50_ values for replicate cultures are shown as dots. e, MDK assay for cefepime-evolved clinical cultures in our evolution protocol (Pb^CEF^) showing the median survivor fraction plotted over time, with error bars representing the median absolute deviation (MAD). Line colors differentiate the parental strain (Pb: black) and evolved cultures that are characteristically tolerant (teal) and persistent (pink). f, heatmap of mutations and their frequencies among individual cultures of Pb^CEF^ and Pb^CEF-R^.

**Supplemental Figure 5.**
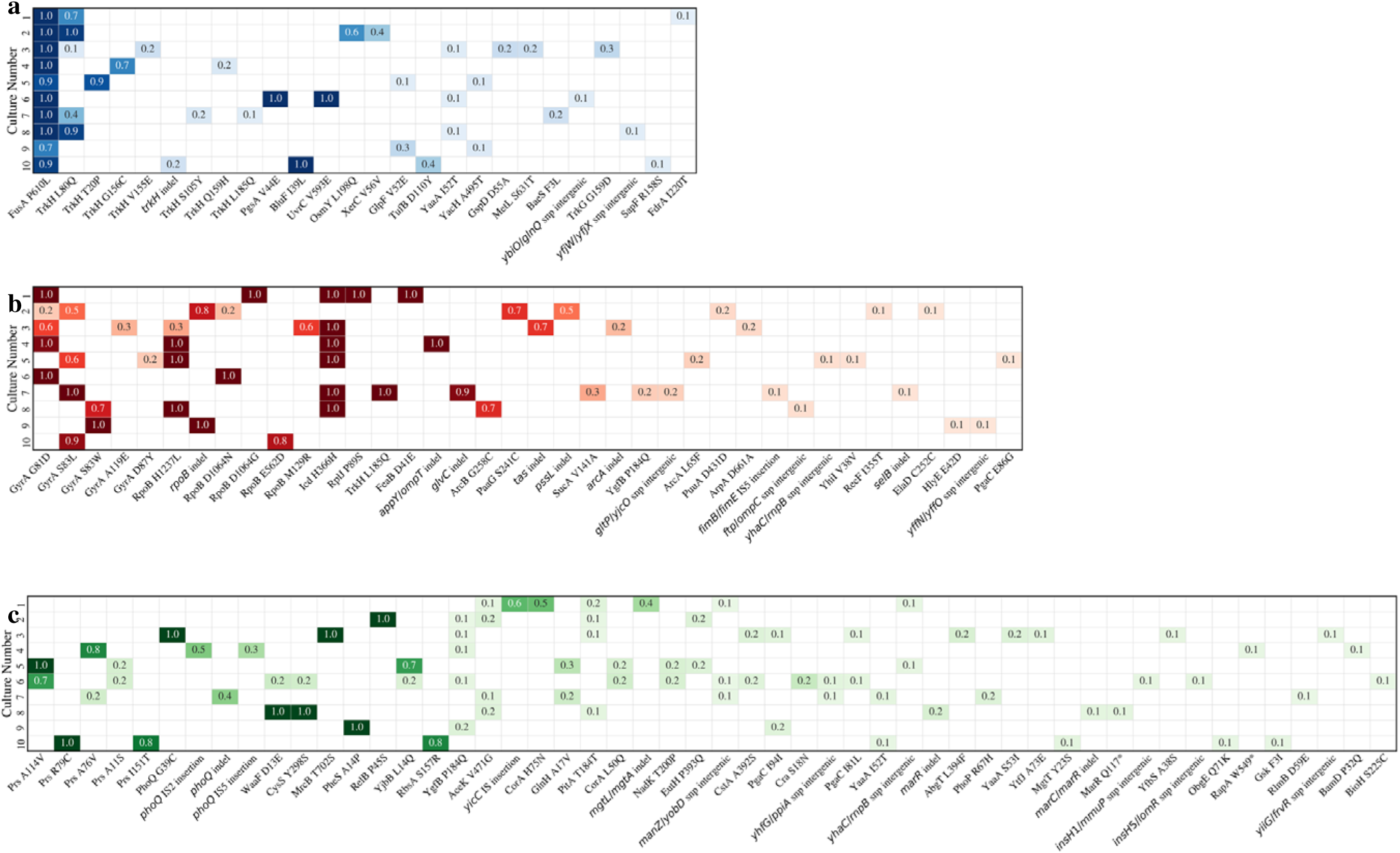
Extended genotypic outcomes of single antibiotic evolutions. Heatmaps of extended mutations and their frequencies among individual cultures: a, MG^AMI^ (blue); b, MG^CEF^ (green); c, MG^LEV^ (red).

**Supplemental Figure 6.**
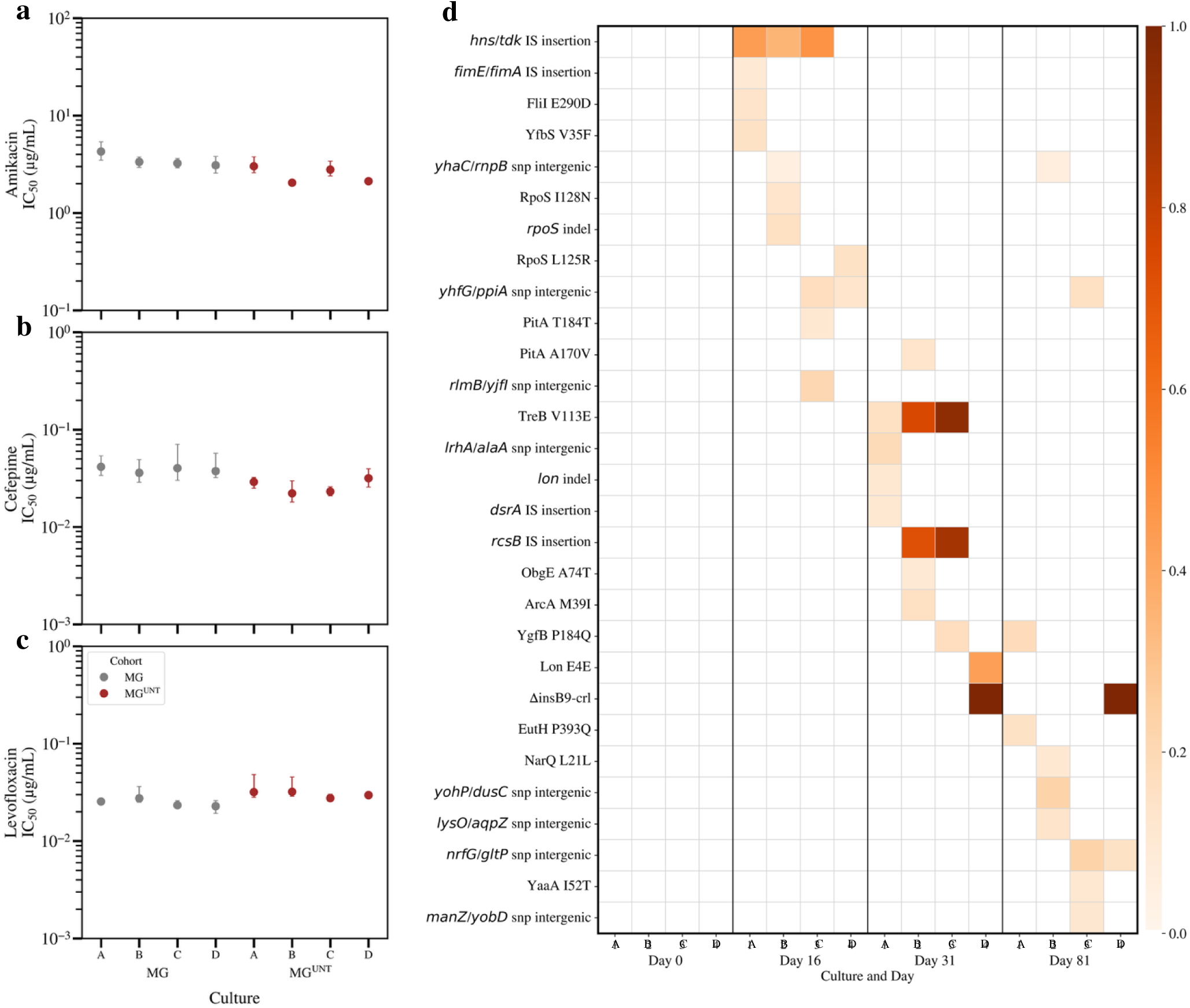
Phenotypic and genetic outcome following untreated passage of MG1655. a-c, IC_50_ values of parent and untreated, passages control populations in each antibiotic, amikacin (a), cefepime (b), levofloxacin (c). d, heatmap of mutational changes throughout untreated passages at key timepoints coinciding with sequential antibiotic treatment.

**Supplemental Figure 7.**
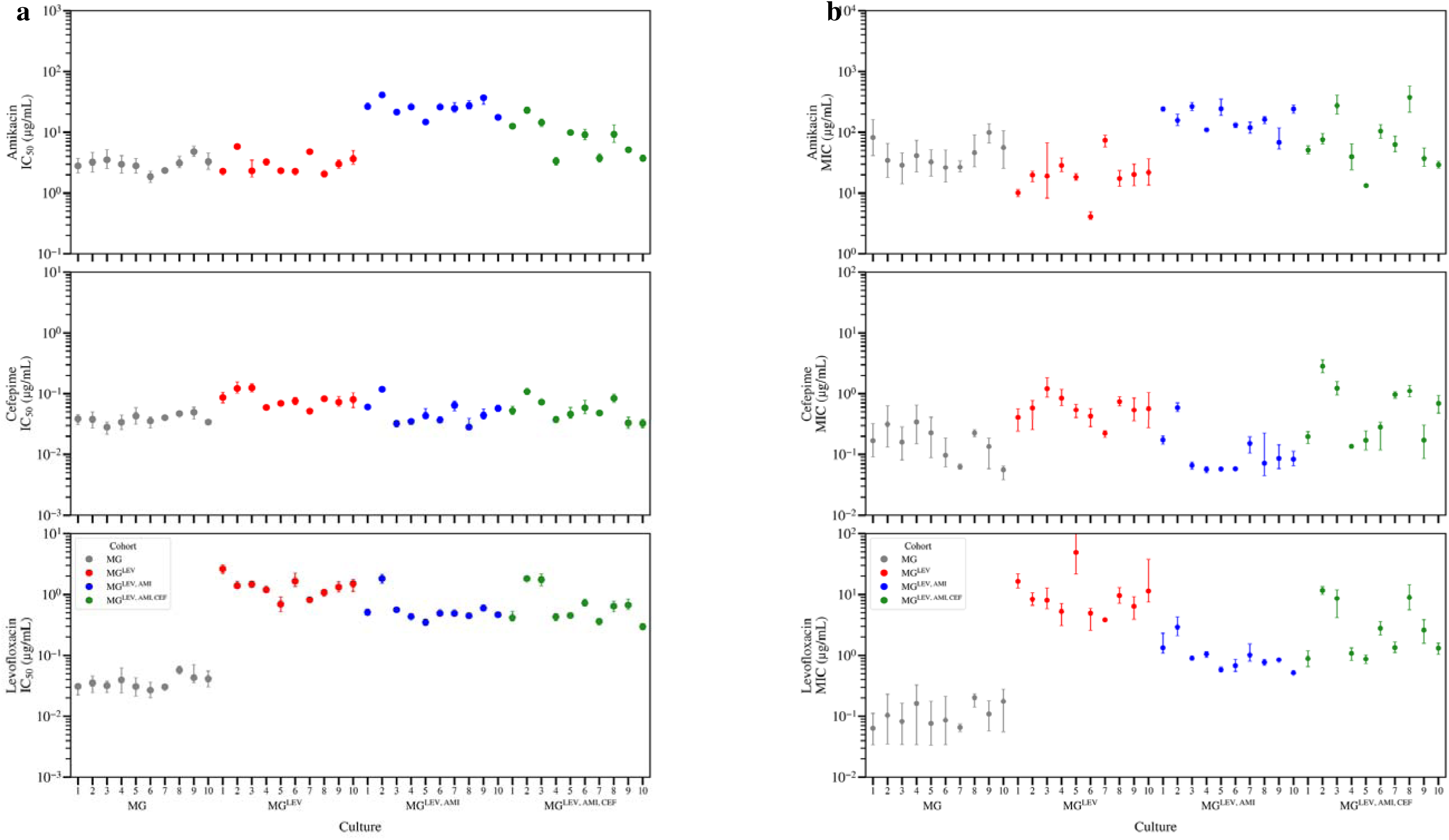
IC50 and MIC values of sequentially evolved cultures. a, Average IC_50_ values (error bars: standard deviation) for individual cultures of parental strain (MG, gray) and cultures sequentially evolved in the presence of levofloxacin (MG^LEV^: red), then amikacin (MG^LEV,AMI^: blue), then cefepime (MG^LEV,AMI,CEF^: green), and b, Average MIC values (error bars: standard deviation) for individual cultures of parental strain (MG, gray) and cultures sequentially evolved in the presence of levofloxacin (MG^LEV^: red), then amikacin (MG^LEV,AM^: blue), then cefepime (MG^LEV,AMI,CEF^: green).

**Supplemental Figure 8.**
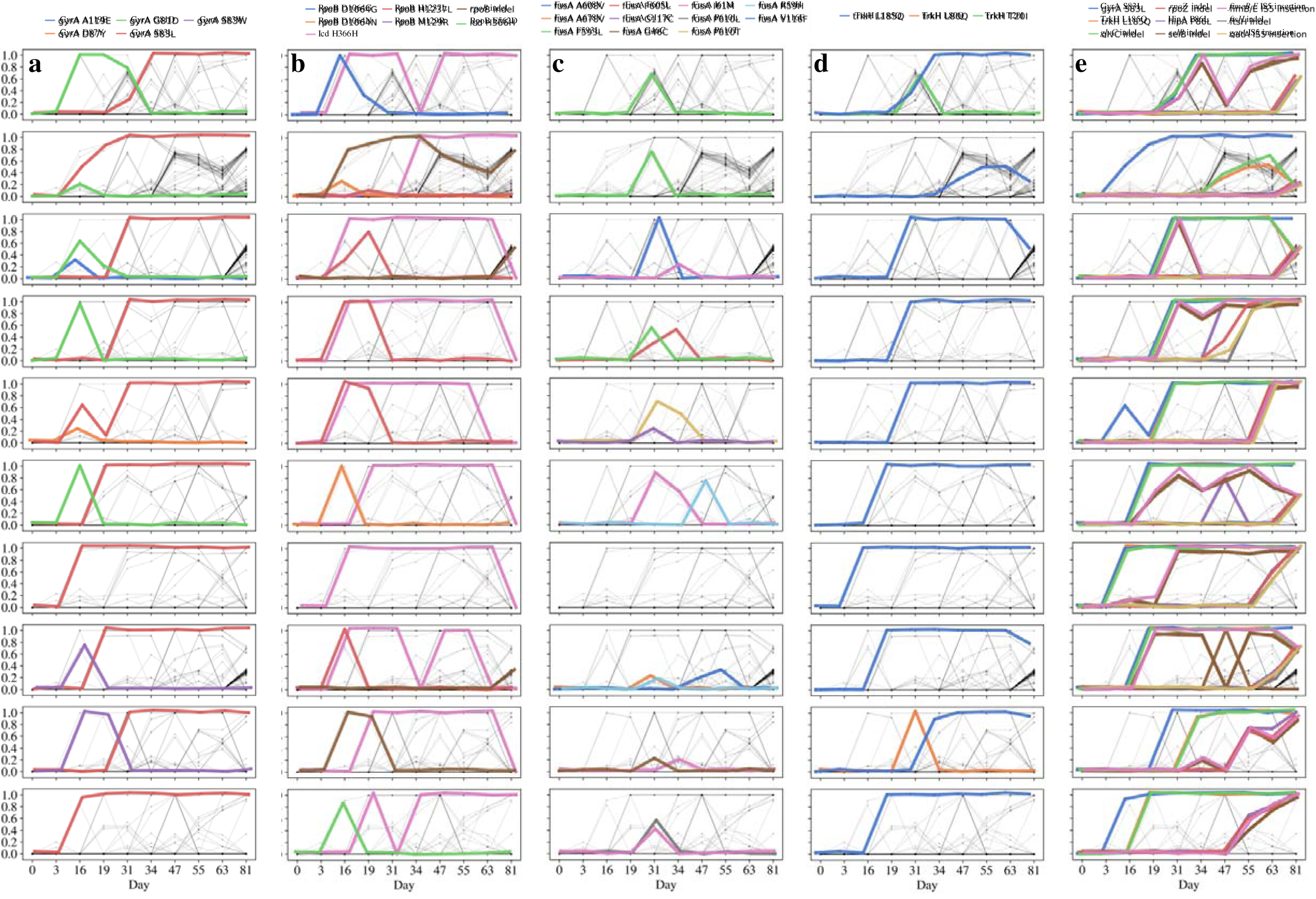
Selected mutation trajectories within each culture. Emergence and fates of selected mutations throughout sequential antibiotic treatment. Columns a-e depict, respectively, mutations of *gyrA*, *rpoB, icd*, *fusA*, *trkH*, and those found to evolve convergently.

**Supplemental Figure 9.**
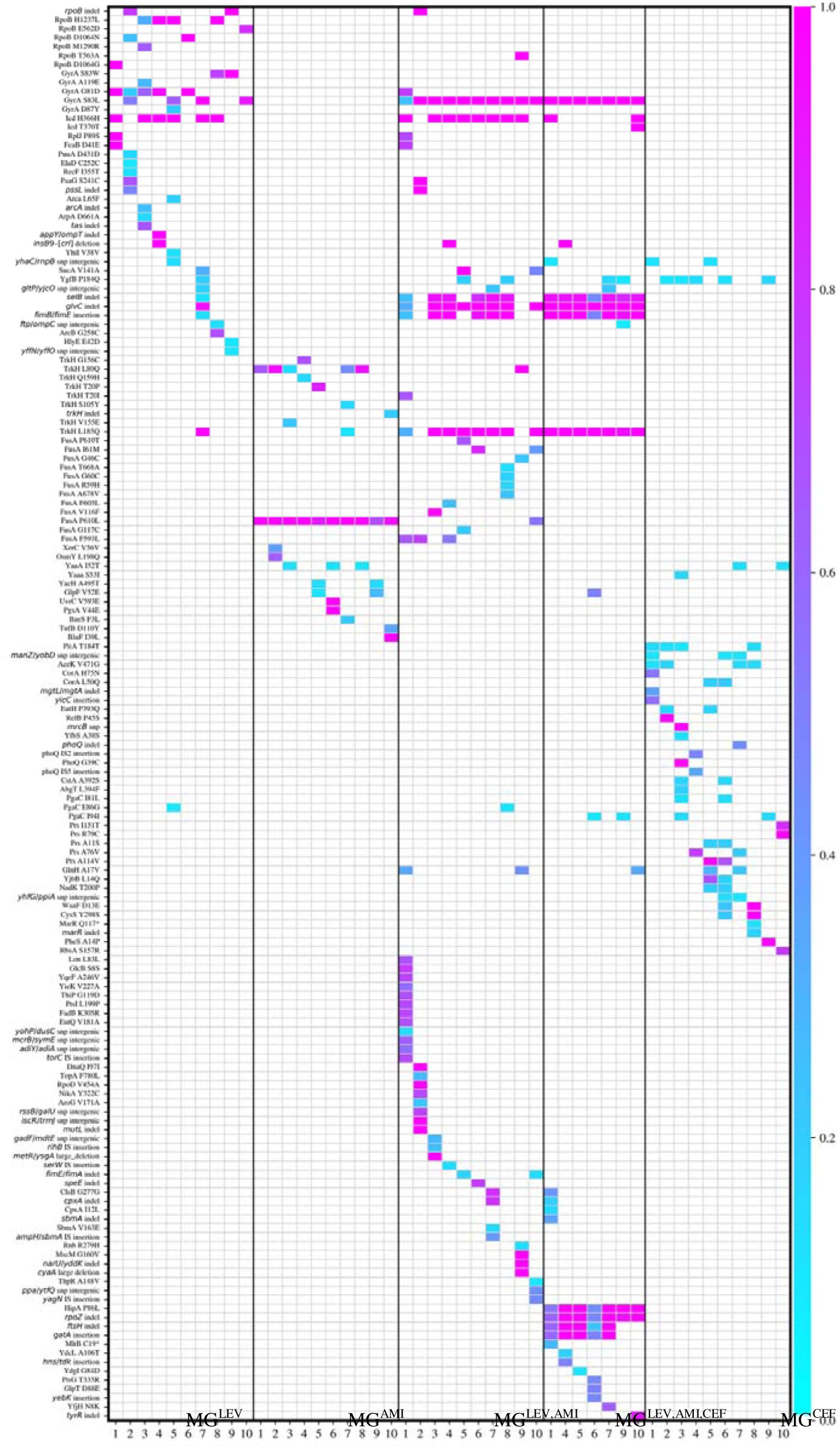
Comprehensive Mutation Frequency Heatmap. All observed mutations among MG^LEV^, MG^AMI^, MG^LEV,AMI^, MG^LEV,AMI,CEF^, MG^CEF^ cohorts (left to right) and their frequencies (color bar, right).

**Supplemental Figure 10.**
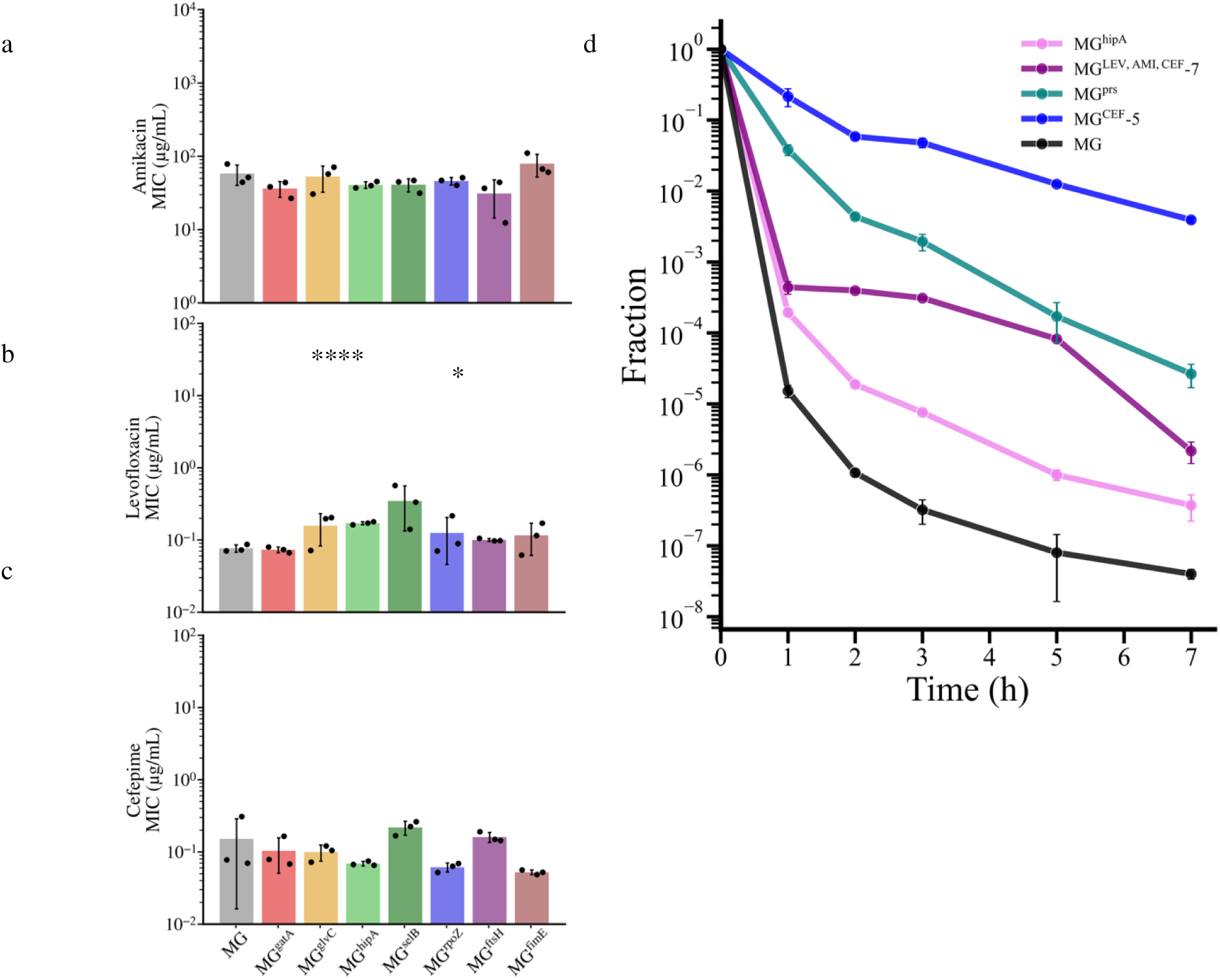
Extended individual mutation characterization. a-c, MIC for individual mutants in each antibiotic, where bar height shows the mean value for three replicates (dots) and error bar indicates standard deviation. d, MDK of mutant strains and evolved cultures with median survival plotted for three replicates and error bar is the median absolute deviation.

**Supplemental Figure 11.**
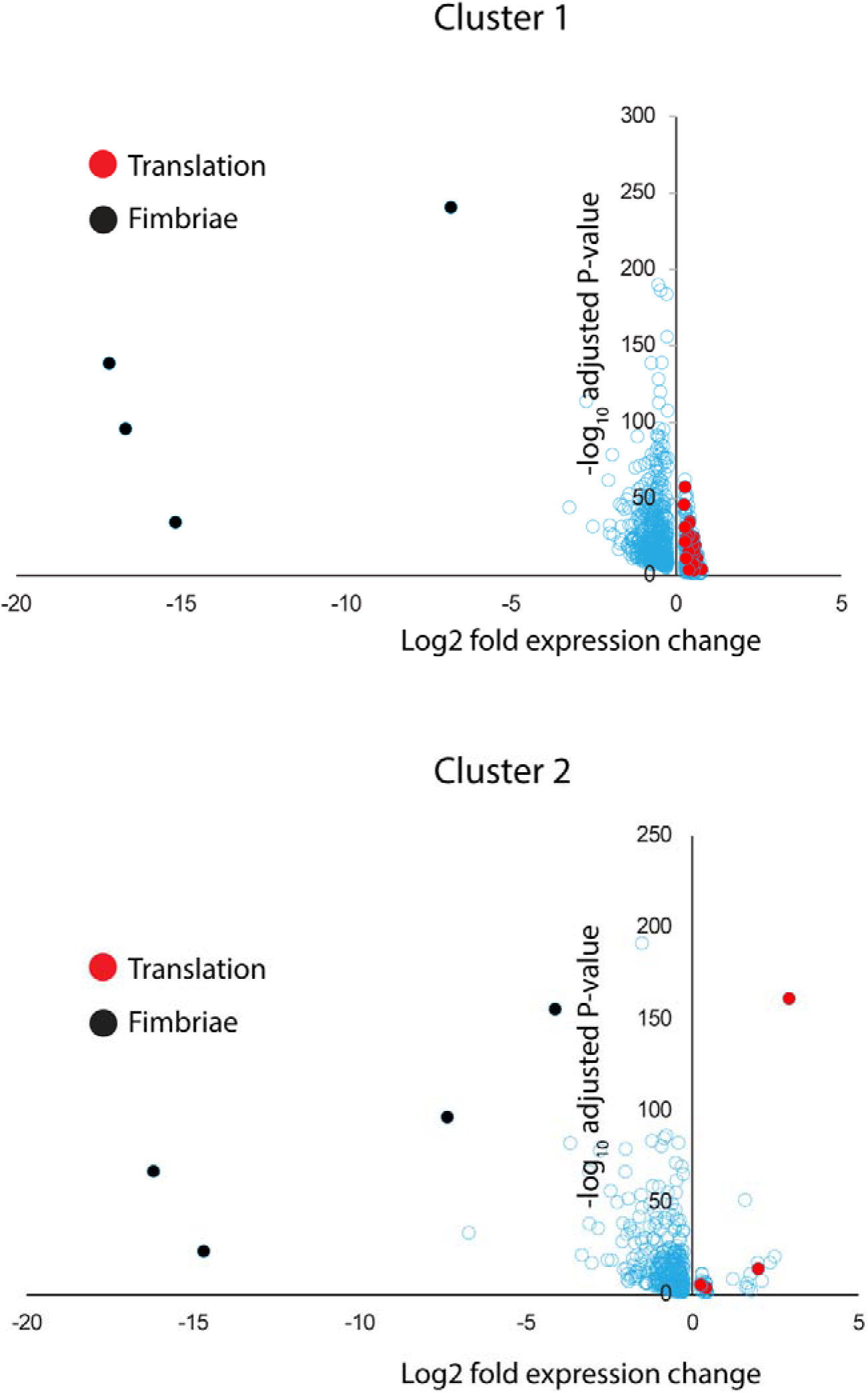
Volcano plots of genes differentially expressed in cluster 1 (top) and cluster 2 (bottom) compared to all other clusters in the combined dataset. Genes involved in protein translation that are overexpressed in these clusters are colored in red. Fimbriae related genes downregulated in these two clusters are colored black. For a full list of differentially regulated genes in each gene-set and P-values please refer to Supplementary Table 3.

**Supplemental Figure 12.**
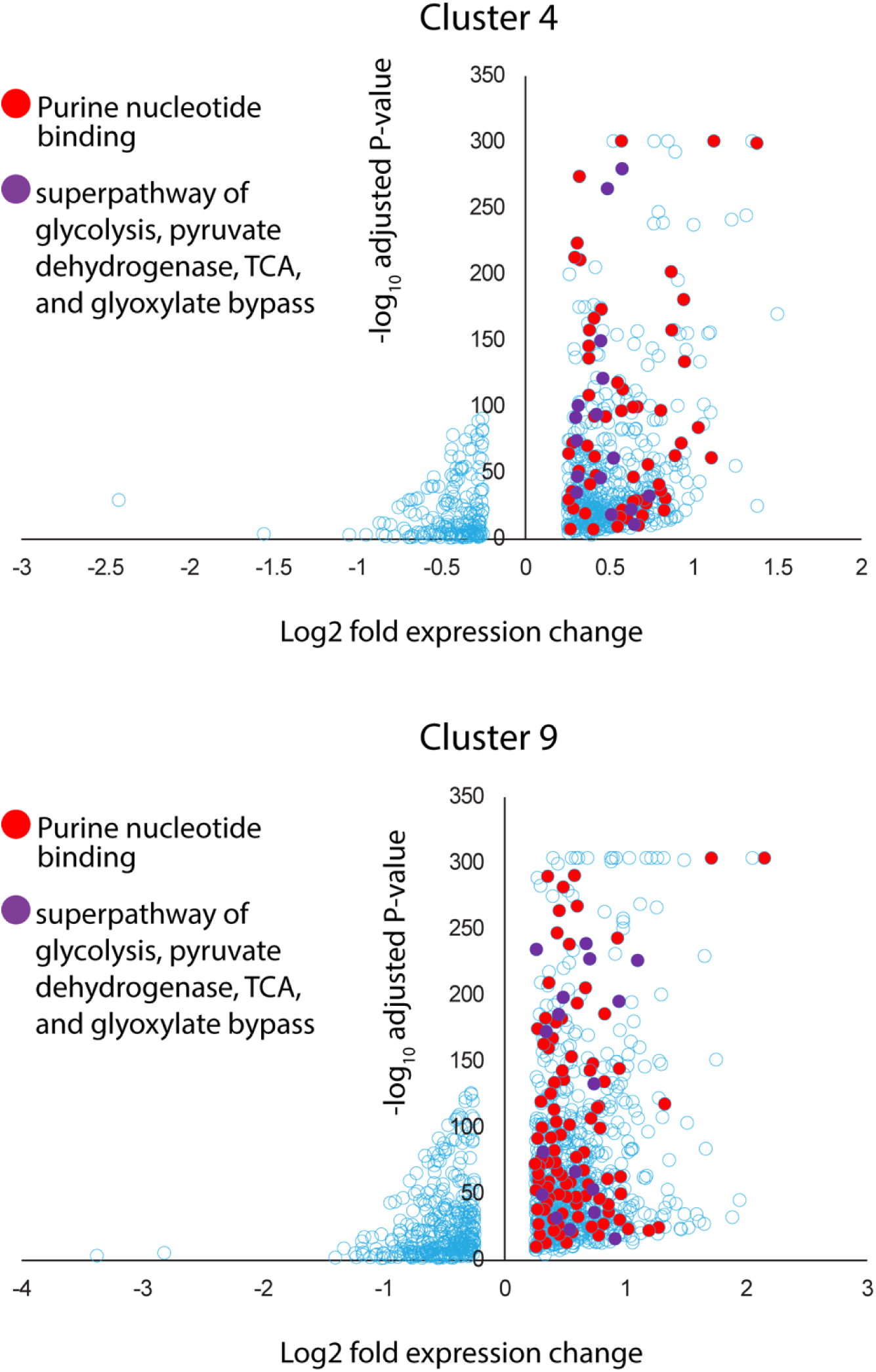
Volcano plots of genes differentially expressed in cluster 4 (top) and cluster 9 (bottom) compared to all other clusters in the combined dataset. Genes involved in purine nucleotide binding that are overexpressed in these clusters are colored in red. Genes in the superpathway of glycolysis, pyruvate dehydrogenase, TCA and Glyoxylate bypass that are overexpressed in these two clusters are colored in purple. For a full list of differentially regulated genes in each gene-set and P-values please refer to Supplementary Table 3.

**Supplementary Table 1.**
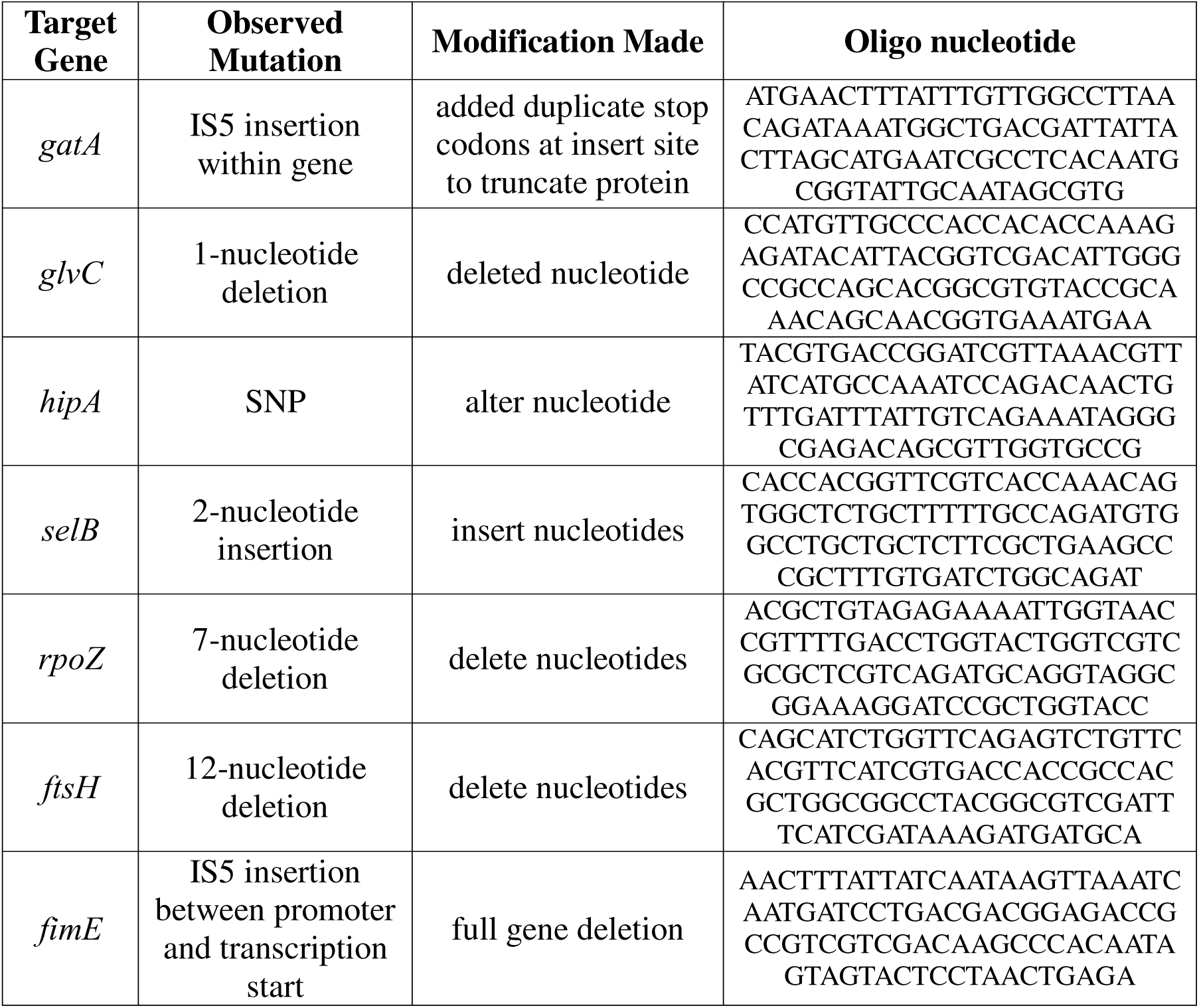
Oligonucleotides for recobineering of individual mutants. Detail of target genes, the mutations observed in each, the approach for modification, and the oligonucleotide used in transformation.

**Supplementary Table 2.**
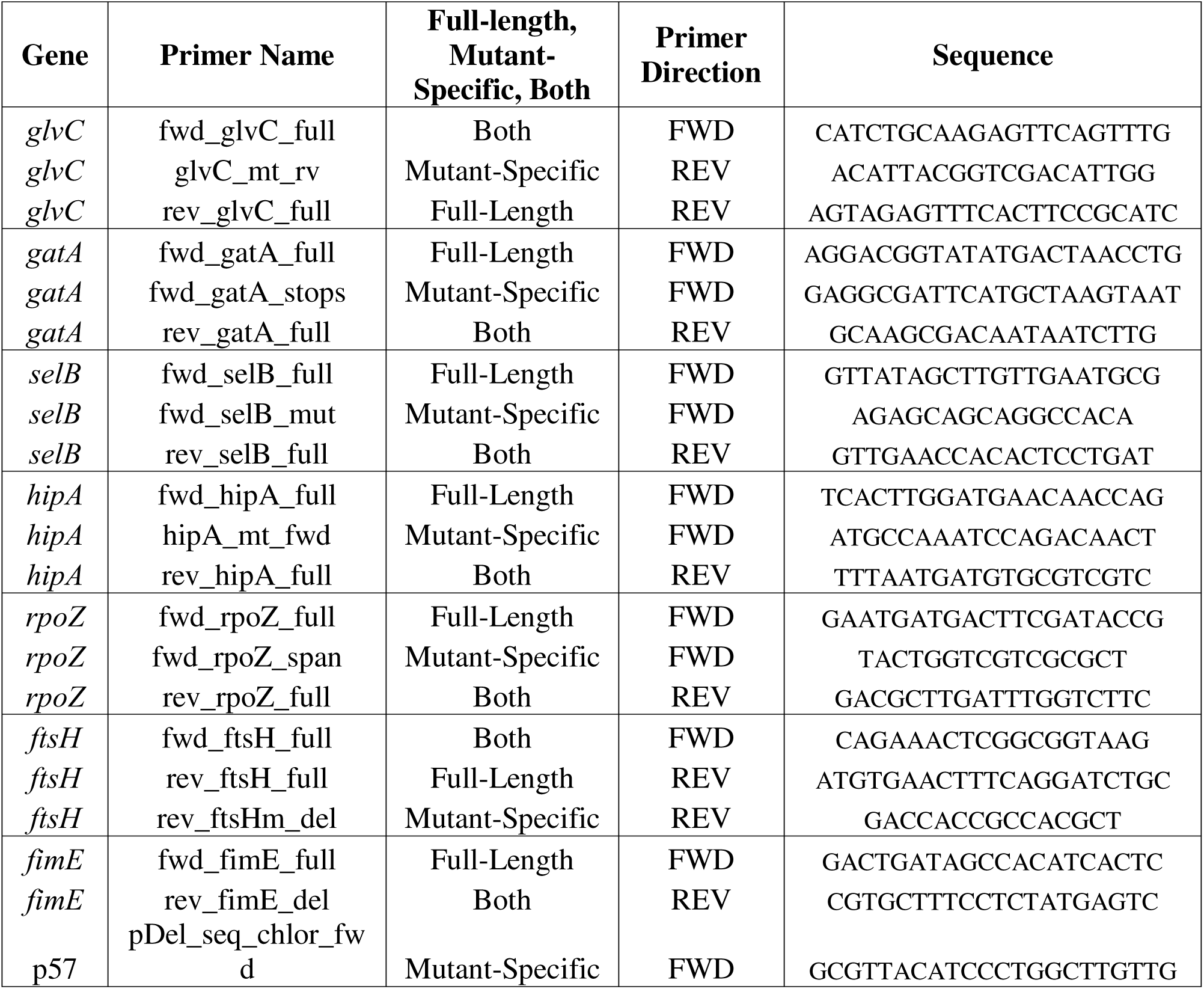
PCR Primers used for screening and confirming oligorecombineering mutations. Details of target PCR products, primer names, product type, primer direction, and sequnce.

**Supplementary Table 3.**
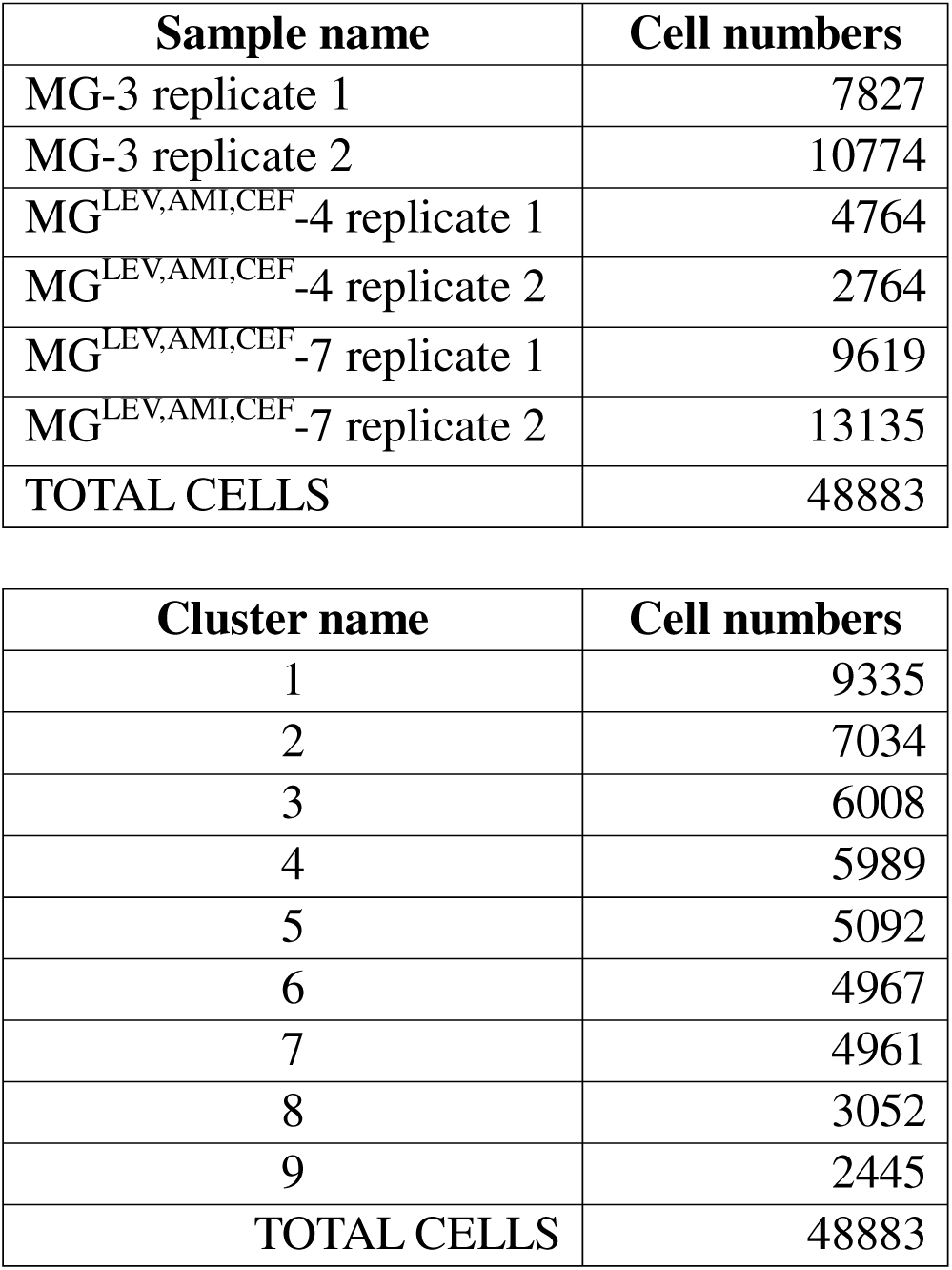
scRNA sequencing samples statistics. Samples (top) and clusters (bottom) and the number of cells representing each group

